# Trypanosomes have divergent kinesin-2 proteins that function differentially in IFT, cell division, and motility

**DOI:** 10.1101/301127

**Authors:** Robert L. Douglas, Brett M. Haltiwanger, Haiming Wu, Robert L. Jeng, Joel Mancuso, W. Zacheus Cande, Matthew D. Welch

## Abstract

*Trypanosoma brucei*, the causative agent of African sleeping sickness, has a flagellum that is crucial for motility, pathogenicity, and viability. In most eukaryotes, the intraflagellar transport (IFT) machinery drives flagellum biogenesis, and anterograde IFT requires kinesin-2 motor proteins. In this study, we investigated the function of the two *T. brucei* kinesin-2 proteins, TbKin2a and TbKin2b, in bloodstream form trypanosomes. We found that compared to other kinesin-2 proteins, TbKin2a and TbKin2b show greater variation in neck, stalk, and tail domain sequences. Both kinesins contributed additively to flagellar lengthening. Surprisingly, silencing TbKin2a inhibited cell proliferation, cytokinesis and motility, whereas silencing TbKin2b did not. TbKin2a was localized on the flagellum and colocalized with IFT components near the basal body, consistent with it performing a role in IFT. TbKin2a was also detected on the flagellar attachment zone, a specialized structure in trypanosome cells that connects the flagellum to the cell body. Our results indicate that kinesin-2 proteins in trypanosomes play conserved roles in IFT and exhibit a specialized localization, emphasizing the evolutionary flexibility of motor protein function in an organism with a large complement of kinesins.

## Introduction

*Trypanosoma brucei spp.* are kinetoplastid parasites that cause African trypanosomiasis (African sleeping sickness) in humans and animals (Brun et al., 2010; Kennedy, 2013). They are transmitted to mammals by the tsetse fly and progress through multiple life-cycle stages including insect vector procyclic form (PCF) and infective mammalian bloodstream forms (BSF). In human African trypanosomiasis (HAT), the bloodstream form proliferates extracellularly in the blood and then in the central nervous system (Kennedy, 2013). HAT is a significant cause of morbidity and mortality in sub-Saharan Africa. However, current treatment options have significant issues with toxicity, cost, difficulty of administration, non-specificity, and drug resistance (Robays et al., 2008; Wilkinson et al., 2008). Thus, the development of new drugs is imperative.

*T. brucei* is highly motile and its motility is driven by a single membrane-bound cilium (flagellum) containing a canonical 9+2 microtubule axoneme with a filamentous paraflagellar rod (PFR) (Langousis and Hill, 2014; Ralston et al., 2009). The flagellum attaches to the cell body along the flagellar attachment zone (FAZ) (Sunter and Gull, 2016; Taylor and Godfrey, 1969), a specialized structure running beneath the plasma membrane along the length of the flagellum. The FAZ contains four specialized subpellicular microtubules (the microtubule quartet, MtQ), associated intracellular membranes contiguous with the endoplasmic reticulum (ER) and nuclear envelope (NE), and a filament system that connects to the axoneme and PFR via several membrane-spanning structures (Kohl and Bastin, 2005; Ralston et al., 2009; Sunter and Gull, 2016; Taylor and Godfrey, 1969). The flagellum beats in distinctive wave patterns that are adapted for movement in the bloodstream (Heddergott et al., 2012; Rodríguez et al., 2009). Movement of BSF *T. brucei* is essential for cell viability and cytokinesis (Broadhead et al., 2006; Ralston and Hill, 2006), development and pathogenesis (Langousis and Hill, 2014), evasion of complement-mediated cell killing (Engstler et al., 2007), and crossing the blood-brain barrier (Kennedy, 2013; Ralston et al., 2009).

The biogenesis of flagella requires the intraflagellar transport (IFT) machinery, which consists of an evolutionarily conserved suite of IFT proteins (Taschner and Lorentzen, 2016; van Dam et al., 2013) and kinesin and dynein motors (Prevo et al., 2017; Scholey, 2013). This machinery is under dynamic regulation to control flagellar structure and function (Heddergott et al., 2012; Ishikawa and Marshall, 2011). The IFT machinery moves cargos along the axoneme in IFT trains (Cole et al., 1993; Cole et al., 1998; Kozminski et al., 1993; Lechtreck, 2015; Pigino et al., 2009) and mediates cargo entry into and exit from the flagellar compartment (Buisson et al., 2013; Dishinger et al., 2010; Verhey et al., 2011). Anterograde IFT transport is driven primarily by kinesin-2 family motor proteins (Scholey, 2013), including two major subfamilies: heterotrimeric kinesin-2, comprising two different kinesin motor subunits (2A and 2B; KRP85 and KRP95) (Cole et al., 1993; Kozminski et al., 1995) and the non-motor kinesin-associated protein (KAP) (Mueller et al., 2005; Wedaman et al., 1996); and homodimeric kinesin-2, which consists of two identical (2C; OSM3) motor subunits (Scholey, 2013; Signor et al., 1999). In metazoans these two subfamilies have been shown to act both cooperatively and independently in IFT (Insinna and Besharse, 2008; Prevo et al., 2015; Snow et al., 2004). Kinesin-2 proteins also participate in extraciliary processes including: cargo transport of organelles, proteins, RNAs, and viruses; mitosis; cytokinesis; cell polarization; and cell adhesion and development (Scholey, 2013).

In this study, we investigate the function of kinesin-2 proteins in BSF *T. brucei.* The *T. brucei* genome encodes approximately 48 kinesins including two kinesin-2 proteins (Berriman et al., 2005; Wickstead et al., 2010b), TbKin2a and TbKin2b. Our bioinformatic analysis suggests that TbKin2a and TbKin2b lack common sequence motifs that are otherwise broadly conserved in kinesin-2 proteins and lack the KAP subunit, which is required for heterotrimer function. We further found that both TbKin2a and TbKin2b contribute to flagellar biosynthesis in BSF *T. brucei.* Interestingly, silencing TbKin2a inhibits cell proliferation, cytokinesis, and motility, whereas silencing TbKin2b does not. Moreover, TbKin2a localizes to the FAZ, suggesting it functions both within the flagellum and cell body. Thus, kinesin-2 proteins in trypanosomes play roles in IFT and may also engage in unanticipated roles.

## Results

### T. brucei has two divergent kinesin-2 proteins

The *T. brucei* genome encodes the kinesin-2 proteins TbKin2a (Tb927.5.2090) and TbKin2b (Tb927.11.13920) (Fig. 1A), which were primarily classified according to their motor domain sequences (Berriman et al., 2005; Wickstead and Gull, 2006; Wickstead et al., 2010b) using phylogenetic inference analysis (Goodson et al., 1994; Wickstead et al., 2010b) (see Fig. S1A for kinesin-2 phylogenetic tree topology from (Wickstead et al., 2010b); see Table S1 for sequence identity comparison of motor domains (MD)). However, sequences in the neck-stalk-tail (NST) region, while not as well conserved as motor domain sequences (average of 17% NST sequence identity versus 54% MD sequence identity), are also important for kinesin function in a variety of metazoan species (De Marco et al., 2001; De Marco et al., 2003; Doodhi et al., 2009; Imanishi et al., 2006; Vukajlovic et al., 2011) (see Table S2 for identity comparison of NST sequences). Nevertheless, systematic analysis of kinesin-2 (or other kinesin) NST sequence motifs in a broad evolutionary context has been limited. We thus sought to further categorize kinesin-2 proteins based on NST sequences.

**Fig. 1:**
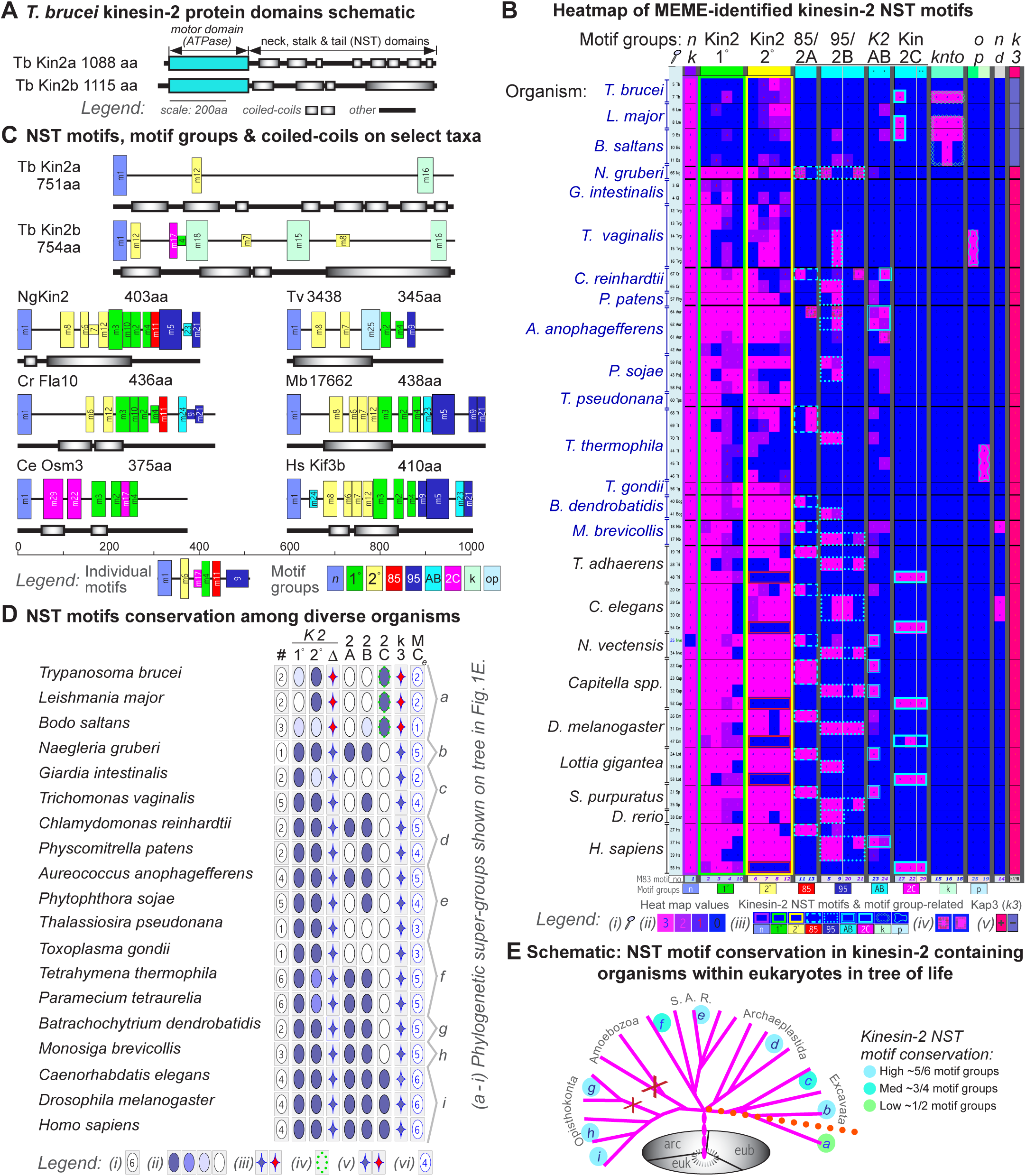
Multiple EM for motif elicitation (MEME) analysis of kinesin 2 sequences from diverse species. **(A)** Schematic illustration of TbKin2a and TbKin2b protein domains. **(B)** Heatmap matrix showing results from a single, representative MEME motif run (MEME 83; out of over 80 runs; see Supplemental File 1 for a list of NST FASTA sequences; see Supplementary File 2 for detailed MEME 83 results). Rows represent 68 taxa (legend (i): species abbreviations require magnification and are explained in Table S3B). Columns represent 26 statistically significant motifs that are clustered into motif groups (based on total E-values; see Fig. S1B for additional information; motif 29 was not statistically significant but is included anyway). Each taxon and motif is color-coded based on motif p-values (legend (ii): from heat map value (hmv) of 3 (magenta) to 0 (blue), indicating taxon-specific individual motif p-values as: 3 = p-value ≤1 × 10^-10^, 2 = p-value >1 × 10^-10^ and ≤1 × 10^-08^, 1 = p-value >1 × 10^-08^ and ≤1 × 10^-05^, and 0 = p-value >1 × 10^-05^). Motif groups are also color-coded (legend (iii): neck (nk or n), motif 1; 1° kinesin-2 (Kin2 1° or 1°), motifs 2, 3, 4, 10; 2° kinesin-2 (Kin2 2° or 2°), motifs 6, 7, 8, 12; KRP85-specific motifs (Kin2A or 2A), motifs 11, 13; KRP95-specific motifs (Kin2B or 2B), motifs 5, 9, 20, 21; present in both KRP85/Kin2A and KRP95/Kin2B (K2AB or AB), motifs 23, 24; Osm3-specific homodimeric (Kin2C or 2C), motifs 17, 22, 29; kinesoplastid-specific (knto or k), motifs 15, 16, 18; other protist-specific (op or p), motifs 19, 25; undefined (nd), motif 14). In rare cases, a single taxon had both 2A and 2B motifs (legend (iv)). The far right column indicates presence/absence of the KAP3 subunit in each taxon (legend (v): with KAP 3 +, red square; no KAP3 -, blue square). **(C)** NST motif and coiled-coil domain schematics for TbKin2a, TbKin2b, and other representative kinesin-2 proteins. Individual motifs are numbered, and each motif sequence is shown in Fig. S1. A key to motif labels and colors is in Fig. S2A. Motif widths are scaled to sequence length, and motif heights indicate p-value ranges (see Fig. S2 (B)). Also see Fig. S2 (D-F) for a more complete set of schematics of representative kinesin-2 proteins. **(D)** NST motif and motif group distribution among a phylogenetically diverse sampling of eukaryotes that have kinesin-2 proteins. Rows represent the listed species. Column designations are indicated under each column as follows: (#) Indicates the number of kinesin-2 taxa in the species (legend (i): number of taxa); (1°, 2°) Indicates the frequency that 1° and 2° motifs occur by color coding (legend (ii): dark blue ≥ 2 motifs at a hmv = 3, medium blue ~1 motif at hmv = 3 or ~2 motifs at hmv = 2, light blue ~1 motif at hmv = 2 or > 2 motifs at hmv = 1, unfilled values < previous categories); (Δ) Indicates whether the order and location of motifs is typical (blue) or atypical (red) (legend (iii): blue star typical, red star atypical). For additional information on motif orders, see Fig. S2C. (2A, 2B) Indicates the frequency that 2A and 2B motifs occur by color coding (legend (ii): dark blue ≥1 motif at hmv = 3, medium blue ≥ ~1 motif at hmv = 2, light blue ≥ ~1 motifs at hmv = 1). (2C) Indicates the presence of a 2C motif assignment (legend (iv): motif assignments to kinetoplastids are outlined with green dots); [H] (k3) Indicates the presence (blue) or absence (red) of genes encoding KAP3 homologs. (legend (v): blue star Kap3 present, red star Kap3 absent) (MC) Indicates the number of kinesin-2 NST motif groups conserved within that species, out of 6 primary motif groups. (legend (vi): number of motif groups). The letters a-i to the right show phylogenetic groups: (a) kinetoplastids, (b) heterobolosea, (c) metamonada, (d) algae and plants that produce flagellated cells, (e) stramenopiles, (f) alveolata (including apicomplexia), (g) fungi (unusual), (h) holozoa, (i) metazoa. **(E)** Schematic based on tree of life presented in (Adl et al., 2012). Species groupings (letters (a-i) from panel (D)) are placed to show distribution within eukaryotic superphyla: ophistokonta (metazoa, holozoa, fungi), amobozoa, SAR (stramenopiles, alveolates and rhizaria), archaeplasitida, and excavata. The degree of motif conservation is color coded. Orange dotted line shows approximate location of possible kinesin-2 divide within the discoba branch of excavates.

We compiled a phylogenetically diverse set of 81 kinesin-2 sequences from 25 species (Table S3A), which represent a broad cross-section of known phylogenetic diversity, and used the multiple Em for motif elicitation (MEME) tool suite to analyze these sequences (Fig. 1B-E; Fig. S1, Fig. S2, Table S3A and S3B). MEME correctly identified the known kinesin-2 neck region sequences that start the NST domain for all kinesin-2 proteins (Case et al., 2000; Vale and Fletterick, 1997). Not unexpectedly MEME identified a single 31 amino acid motif consistent with a kinesin-2 neck domain at the beginning of the NST domain for all kinesin-2 proteins in our data set (Case et al., 2000; Vale and Fletterick, 1997). MEME also identified over 20 additional, statistically significant NST sequence motifs that can be sorted into distinct ordered motif groups that are broadly conserved across all kinesin-2 containing eukaryotic superfamilies from Metazoa to Excavata (Fig. 1D, E). Notably, motifs forming a primary (1°) motif group were found to be common to almost all protist and metazoan taxa. A secondary (2°) motif group that is frequently associated with coiled-coil domains, as well as motifs that are specific to subgroups 2A and/or 2B, were also widely shared among protists and metazoa (Fig. 1B-E; Fig. S1B, C; Fig. S2). The assignment of protist kinesin-2 taxa into subgroups 2A and 2B was not observed in the phylogenetic analyses based on motor domain sequences. MEME also assigned 2C motifs to metazoan kinesin-2C taxa, but such motifs were absent from protist taxa. The primary exception was a single 2C motif shared in kinetoplastids and metazoans, as discussed below (Fig. 1B-E; Fig. S1, Fig. S2).

The greatest divergence in NST sequence motifs occurs within the kinetoplastids including *T. brucei* (Fig. 1B-E). Kinetoplastid kinesin-2 NST domains are longer, have more extensive predicted coiled-coil sequences, and lack most motif groups that are present in other protists and metazoans (Fig. 1; Fig. S2). Instead, the kinetoplastids have unique motif groups (knto/k) that are specific to these organisms. Parasititic kinetoplastid species (Julkowska and Bastin, 2009) and the free living *B. saltans*, lack the KAP subunit (k3) (Fig. 1B, D), which is required for kinesin-2 heterotrimer function and is present in all other organisms included in our analysis. Surprisingly, MEME assigned to the kinetoplastid homologs of TbKin2b a single, statistically significant kinesin-2C (OSM3) motif that is also present only in metazoan taxa (Fig. 1B-E; Fig. S1, Fig. S2). Together, these results suggest that kinetoplastid kinesin-2 proteins form a unique subfamily and are unlikely to have a heterotrimeric structure consisting of 2A, 2B and KAP subunits. Thus, MEME analysis of NST sequences suggests that kinesin-2 proteins in *T. brucei* have diverged from such proteins in other organisms, raising questions about the extent to which canonical functions of these proteins are conserved in trypanosomes, and if novel functions have evolved.

### TbKin2a distributes along the length of the flagellum, with stronger localization near the basal body

To assess the function of kinesin-2 proteins in *T. brucei*, we first raised antibodies that specifically recognize the ~130 kD TbKin2a protein on immunoblots of whole-cell lysates (supplementary material Fig. S3) and assessed TbKin2a localization in BSF cells by immunofluorescence microscopy. TbKin2a localized in a punctate pattern along the flagellum (Fig. 2A). During the cell cycle, TbKin2a staining intensity on flagella increased and relative staining between old and new flagella remained roughly equivalent, except that TbKin2a typically showed enhanced staining on new flagella relative to old flagella primarily during the 1N1K stage. This reflects the cytoskeletal changes associated with the emergence of the new flagellum (Gluenz et al., 2011; Ikeda and de Graffenried, 2012; Lacomble et al., 2010). We also determined the mean fluorescence intensity along full-length flagella. Using the peak intensity of DAPI-stained kinetoplast DNA as a posterior reference point (Fig. 2B), we determined that the flagellar intensity profile (on both old and new flagella) had two peaks at ~ 0.7 μm and ~ 2.1 μm, approximating locations near the basal body and flagellar collar. Past the second peak, staining diminished to the anterior tip, where it increased slightly, with 65% of integrated staining intensity being confined to the proximal half of the flagellum.

**Fig. 2:**
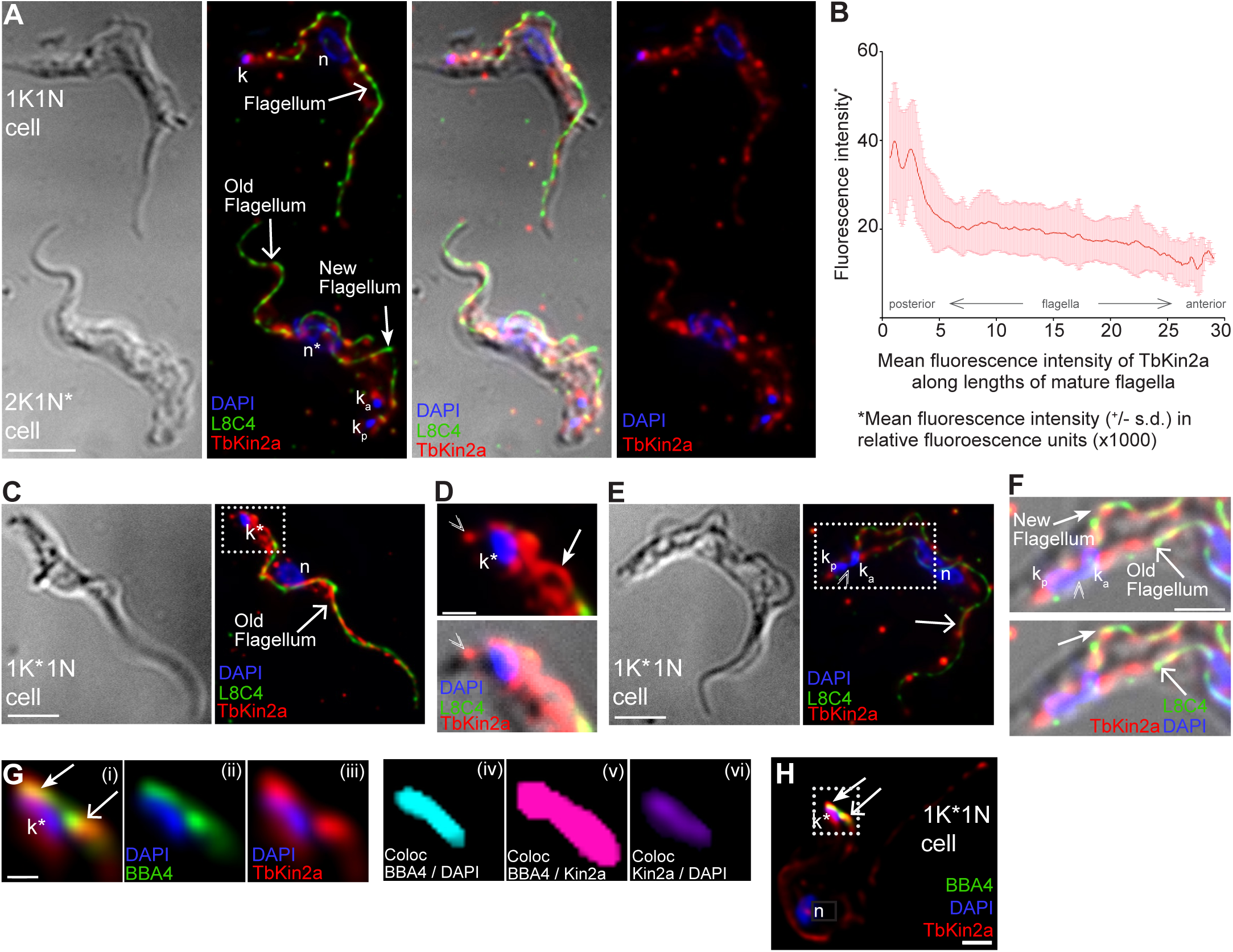
TbKin2a localizes to flagella and basal bodies. **(A)** BSF (90-13) cells F/M-fixed and stained for TbKin2a (red), paraflagellar rod protein (PFR) as a flagellar marker (green, L8C4) and DNA (blue, DAPI). Bar = 5 μm. **(B)** TbKin2a mean relative fluorescence intensity measured along individual M-fixed flagella mature BSF flagella (measured every 0.1 μm from posterior to anterior ends, starting adjacent to kinetoplast near proximal end of mature basal body). Mean flagellum length = 17 μm. Error bars = SD (n = 139). **(C)** Images of a F/M-fixed 1K^∗^1N cell stained for the PFR (green, L8C4), TbKin2a (red), and DNA (blue, DAPI). Bar = 3 μm. **(D)** Magnified area boxed in (C). Double V indicates unusual posterior end cell staining by TbKin2A that was typical only in this stage of cell cycle. Bar = 1 μm. **(E)** Images of F/M-fixed 1K^∗^1N cell stained for the PFR (green, L8C4), TbK2a (red) and DNA (blue, DAPI). Bar = 3 μm. **(A-E)** Standardized abbreviations and symbols include: Fixation method, M-fixed = methanol fixed, F-fixed = formaldehyde fixed, F/M-fixed = sequential formaldehyde then methanol fixed; images are differential interference contrast (DIC) or fluorescence corresponding to two-dimensional (2D) projections from a three-dimensional (3D) deconvolved stack; wide arrow = flagellum (interphase) or old flagellum; barbed arrow = new flagellum; k = kinetoplast; k^∗^ = dividing kinetoplast; k_a_ = anterior kinetoplast; k_p_ = posterior kinetoplast; n^∗^ = mitotic nucleus; n = nucleus. **(F)** Magnified area boxed in (E). Doubled V indicates narrow area connecting two parts of nearly fully divided kinetoplast. Bar = 1.5 μm. **(G)** Magnified area boxed in (H) showing: (i-iii) Region near the basal bodies in wild-type 1K^∗^1N cell stained for basal bodies (green, BBA4), TbKin2a (red) and kinetoplast DNA (blue, DAPI); (iv-vi) Colocalization of threedimensional voxels between BBA4 and TbKin2a (pink), DNA and TbKin2a (purple), and DNA and BBA4 (blue). Bar = 0.75 μm. **(H)** Image of a 1K^∗^1N cell from which panels in (G) are taken. Bar = 2.4 μm.

TbKin2a also localized in the region of the kinetoplasts and basal bodies throughout their duplication cycle (Fig. 2C-E). There was extensive overlap between TbKin2a and basal bodies, as marked by the proximal-end basal body marker, BBA4 (Dilbeck et al., 1999; Woods et al., 1989) (Fig. 2E; threshold-adjusted Mander’s colocalization coefficient for BBA4/TbKin2a of 0.738 (Manders et al., 1993)). Moreover, TbKin2a and BBA4 colocalized with kinetoplastid DNA near the proximal ends of basal bodies (Fig. 2E). Thus, TbKin2a is positioned on the basal body adjacent to the kinetoplast. Given its localization pattern, TbKin2a may be a useful marker to visualize progressive changes in the positioning and orientation of kinetoplasts, basal bodies, and flagella during cell-cycle progression and new flagellum biogenesis.

### TbKin2a colocalizes with IFT proteins primarily at the flagellum base

Although the canonical role of kinesin-2 proteins is in IFT (Lechtreck, 2015; Scholey, 2013), the divergent sequence of TbKin2a raised the question of whether it retains its function in IFT in trypanosomes. In *T. brucei*, full complements of IFT complex B (anterograde; IFTB) and IFT complex A (retrograde; IFTA) proteins have been identified and localized to the flagellum and basal body regions (Absalon et al., 2008; Adhiambo et al., 2009; Franklin and Ullu, 2010), and shown to function in IFT including the movement of cargo trains (Blisnick et al., 2014; Buisson et al., 2013; Huet et al., 2014). We co-stained for TbKin2a with IFTB marker IFT172 (Fig. 3A-D) or IFTA marker IFT144 (Fig. 3E). As previously reported in PCF cells (Absalon et al., 2008), in BSF cells these markers exhibited punctate staining along the flagellum and increased intensity near basal bodies. Along the flagellum, there was little colocalization between TbKin2a and either IFT172 or IFT144 (Fig. 3A, C, E, F; mean Mander’s colocalization coefficients 0.164 and 0.278, respectively). However, in the region near the kinetoplasts, basal bodies, and proximal ends of flagella, colocalization between TbKin2a and each IFT protein was observed (Fig. 3B, D, E, F; mean Mander’s colocalization coefficients of 0.655 and 0.636, for IFT172 and IFT144, respectively). This suggests that despite its sequence divergence, TbKin2a interacts with IFT complex proteins, primarily near the flagellum base.

**Fig. 3:**
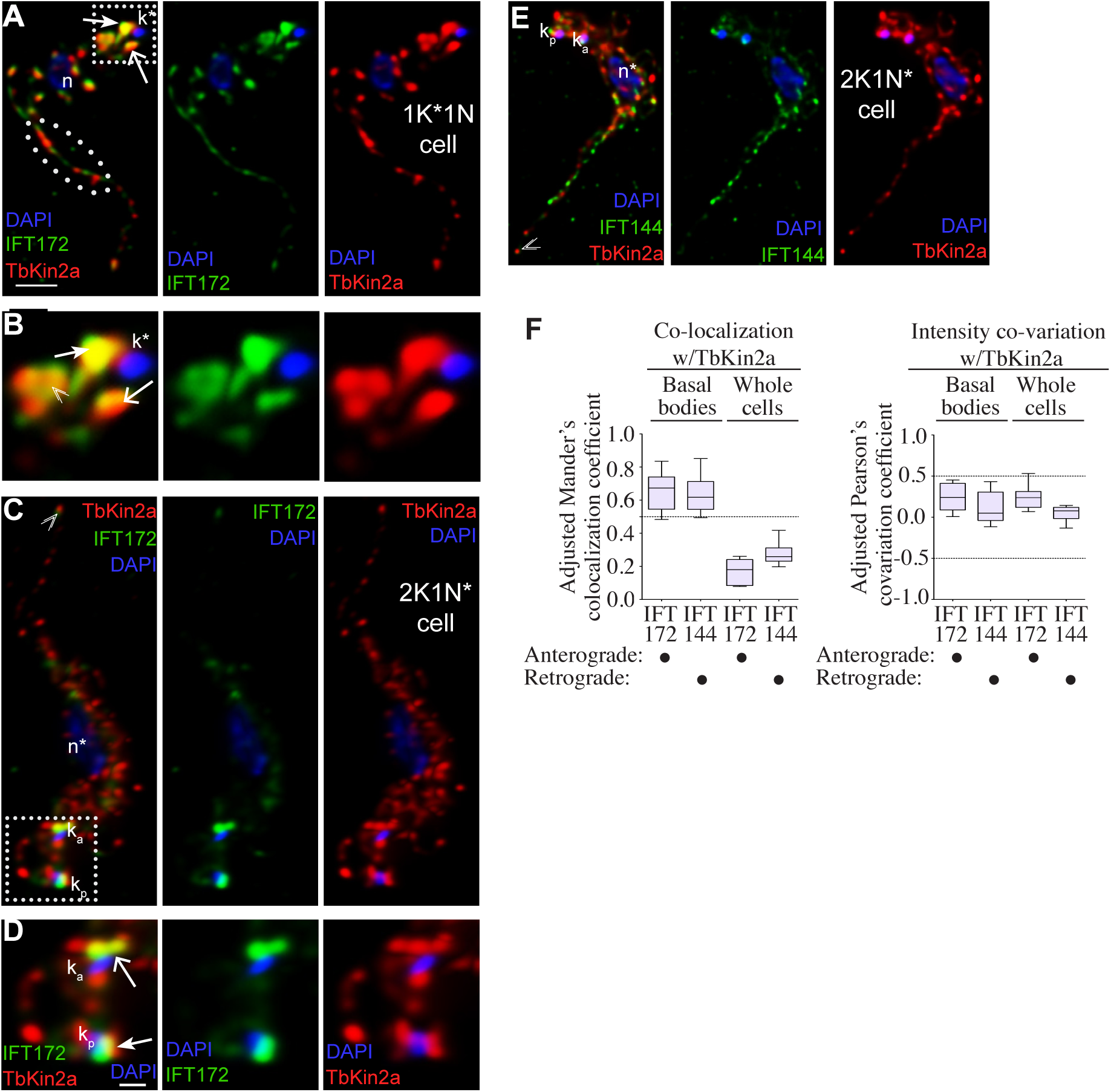
Colocalization of TbKin2a with IFTA and IFTB proteins. **(A)** 1K^∗^1N cell stained for IFT anterograde complex B protein IFT172 (green), TbKin2a (red), and DNA (blue, DAPI). Voxels with colocalized IFT172 and TbKin2a are gold. Dotted-ellipse shows area in anterior half of flagellum having almost no colocalization. Bar = 2 μm. **(B)** Magnified of boxed area in (A) showing the region near basal bodies. Bar = 0.7 μm. See also comment in (D) **(C)** 2K1N^∗^ cell stained as in (A). Bar = 4 μm. **(D)** Higher magnification view of boxed area in (C) showing the region near basal bodies. Bar = 0.7 μm. Comparison of (B) and (D) staining near basal bodies shows typically increased TbKin2a and IFT172 staining at and near base of new flagellum (double-V symbol) for 1K^∗^1N stage cells in (B) relative to 2K1N^∗^ cells in (D). **(E)** 2K1N^∗^ cell stained for retrograde complex A protein IFT144 (green), TbKin2a (red) and DNA (blue, DAPI). Bar = 2 μm. Note comparative staining for retrograde Complex A protein IFT144 and TbKin2a in (E) is generally similar to that between anterograde Complex B protein IFT172 and TbKin2a in (C). **(A-E)** All images are of cells that are F/M-fixed and stained as labeled, presented as 2D projections from a 3D deconvolved stack. Standardized abbreviations and symbols have the same meaning as in Fig. 2. **(F)** Mander’s colocalization coefficient and Pearson’s covariation coefficient for the region surrounding the basal bodies, and the whole cell, to quantify colocalization of TbKin2a with IFT172 or IFT144 (for IFT172, n = 11; for IFT144 n = 7).

### TbKin2a localizes to the FAZ throughout the cell cycle

Given the tight association of the flagellum with the FAZ, we sought to distinguish whether TbKin2a also localizes to the FAZ. The flagellum can be detached from FAZ and cell body by detergent extraction, which permits separate visualization of the components localized to each structure (Robinson et al., 1991). Surprisingly, at all stages of the cell cycle, TbKin2a colocalized with the FAZ marker L3B2 (Kohl et al., 1999) in a continuous punctate pattern from the initiation of the FAZ (start of L3B2 staining) near the flagellar collar (Lacomble et al., 2010; Lacomble et al., 2009) to the anterior tip (Fig. 4A) (in contrast, we did not observe evidence of IFT172 colocalization with FAZ, not shown). Because detergent extraction significantly diminished TbKin2a staining on flagella (also observed with IFT proteins) (Absalon et al., 2008), we sought to identify conditions to separate the flagellum and FAZ without detergent extraction following fixation. We found that the flagellum and FAZ were sometimes partially separated due to shear forces during mounting. Under these circumstances, TbKin2a could always be seen on both structures (Fig. 4B, C). We found that we could also use deliberate but controlled shearing to induce similar separation of the flagellum and FAZ post-fixation and that under these circumstances, TbKin2a could also always be seen on both structures (Fig. 4D, E). Collectively, these results indicate that TbKin2a associates with the FAZ as well as with the flagellum.

**Fig. 4:**
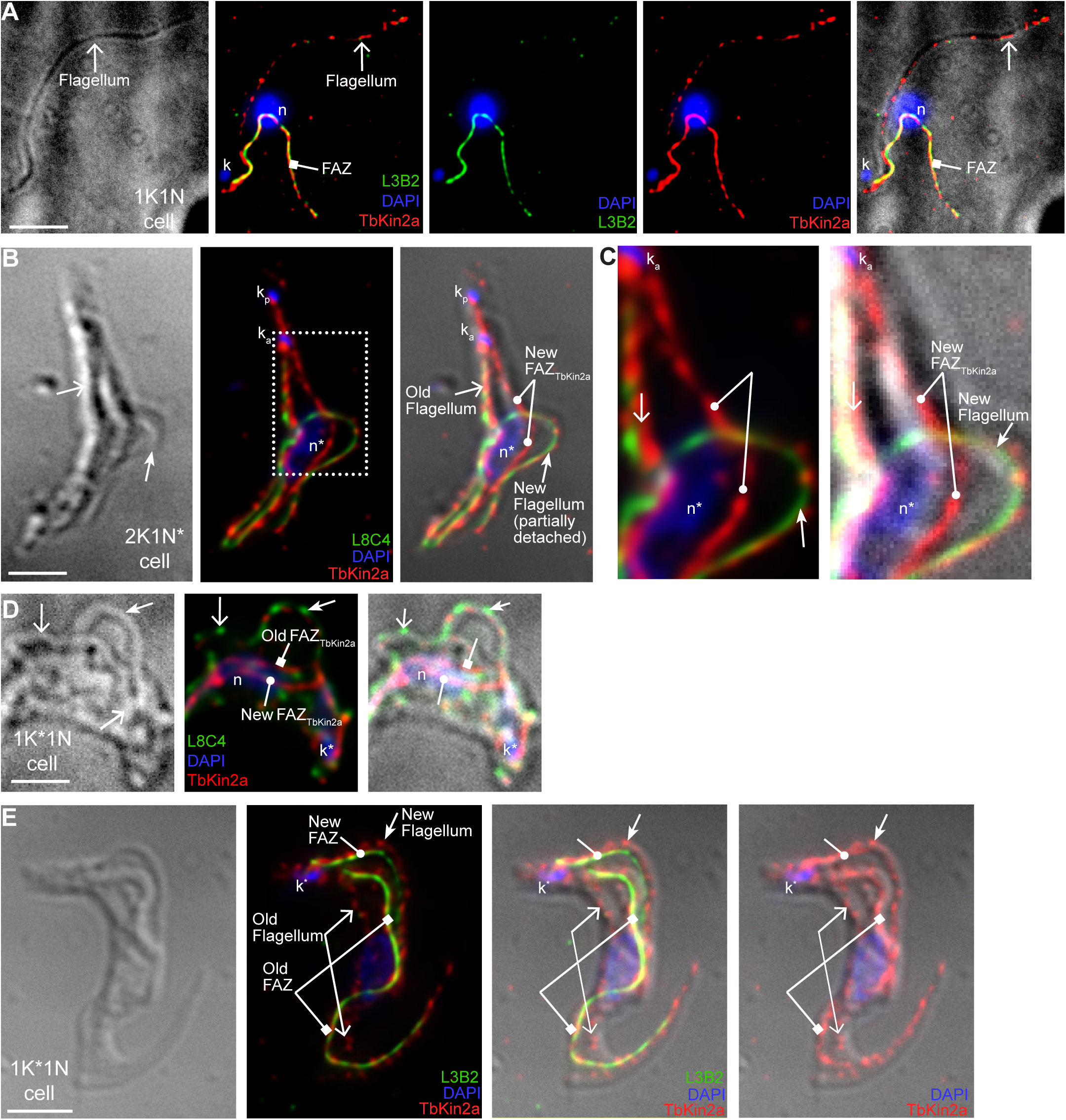
TbKin2a localizes to the FAZ. **(A)** 1K1N detergent-extracted cell stained for TbKin2a (red), FAZ marker protein FAZ1 (green, L3B2) and DNA (blue, DAPI) showing TbKin2a staining on both flagellum and FAZ. Note that detergent extraction protocol included additions of ATP and Mg^2+^, see Materials and Methods. Bar = 2.5 μm. **(B)** 2K1N^∗^ cell, subjected to shear forces post-fixation such that the new flagellum has become partially detached and separated from the new FAZ; stained for TbKin2a (red), flagellar PFR (green, L8C4) and DNA (blue, DAPI). **(C)** Magnified image of boxed area from (B) showing independent TbKin2a staining on the FAZ as well as on the detached flagellum colocalized with the PFR. **(D)** 1K^∗^1N cell, subjected to defined shear forces post-fixation such that both old and new flagella are partially detached (see Materials and Methods), stained with TbKin2a (red), flagellar PFR (green, L8C4), and DNA (blue, DAPI). **(E)** 1K^∗^1N cell subjected to defined shear forces post-fixation such that the old flagellum is now separated from the old FAZ while the new flagellum and new FAZ are still unseparated; stained with TbKin2a (red), FAZ1 marker (green, L3B2), and DNA (blue, DAPI). **(A-E)** All images are of cells that are F/M-fixed and are taken using DIC microscopy, or fluorescence microscopy corresponding to 2D projections from a 3D deconvolved stack. Standardized abbreviations and symbols are the same as those in Fig. 2., and new symbols include the following: bar with filled square = old FAZ; bar with filled circle = new FAZ.

### Silencing of TbKin2a inhibits growth of BSF T. brucei, whereas silencing of TbKin2b does not

To determine the importance of kinesin-2 proteins in BSF *T. brucei*, we used RNA interference (RNAi) to silence expression of TbKin2a, TbKin2b, or both. TbKin2a mRNA was diminished at 24 h and 48 hpi post induction (hpi) of dsRNA expression in TbKin2a-silenced cells, but remained unaffected in TbKin2b-silenced cells, as assessed by RT-PCR (Fig. 5A) and qRT-PCR (Fig. 5B). Similarly, TbKin2b mRNA was diminished at 24 hpi and 48 hpi in TbKin2b-silenced cells but was unaffected in TbKin2a-silenced cells (Fig. 5A, B). Both mRNAs were diminished in TbKin2a/2b-silenced cells. For TbKin2a, protein expression was strongly diminished after 48 hpi (Fig. 5A).

**Fig. 5:**
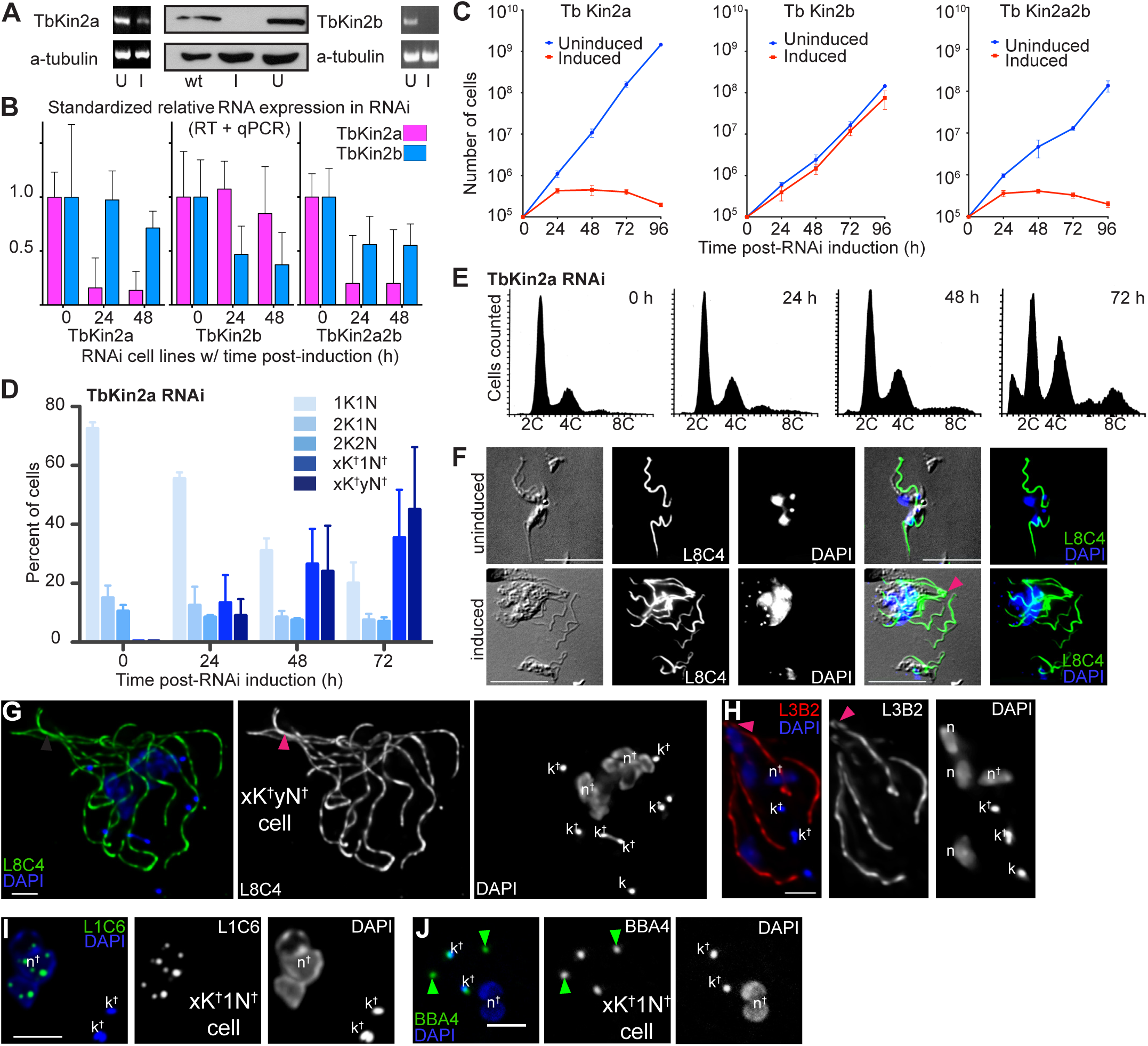
Effect of RNAi silencing of TbKin2a and TbKin2b on cell proliferation and division. **(A)** Left and right: TbKin2a and TbKin2b mRNA levels in RNAi-uninduced (U) and RNAi-induced (I) cells at 48 hpi by RT-PCR, with α-tubulin mRNA levels as a control. Center: TbKin2a protein levels in wild type (wt), RNAi-induced (I), and uninduced (U) cells at 48 hpi by Western blotting using anti-TbKin2a (top) or anti-tyrosinated-α-tubulin YL1/2 (control). **(B)** Comparison of uninduced versus induced normalized relative TbKin2a and TbKin2b mRNA levels in TbKin2a-silenced (n = 4), TbKin2b-silenced (n = 2), and TbKin2a/2b-silenced cells (n = 4), as measured by qRT-PCR. Error bars = SD. Relative gene expression and SD was determined using the 2^−ΔΔC^T method. **(C)** Cell number versus hpi for cells uninduced or induced for RNAi to silence TbKin2a, TbKin2b, or TbKin2a/2b. Data is from two independent biological experiments, each with three technical replicates. Error bars = SD. **(D)** Cell morphology phenotypes with nucleus (N) and kinetoplast (K) counts for TbKin2a RNAi uninduced (n = 197 cells) and induced (n = 230 cells) 0-72 hpi. 1†N, single abnormally large nucleus. **(E)** FACS plots of DNA content for TbKin2a RNAi cells at 0-72 hpi. Error bars = SD. **(F)** Uninduced (top) or induced (bottom) M-fixed cells at 72 hpi, stained for the PFR (green, L8C4) or DNA (blue, DAPI). Bar = 10 μm. **(G)** TbKin2a-silenced cells at 48 hpi stained as in (F). Bar = 5 μm. **(H)** TbKin2a-silenced cell that was detergent-extracted and stained for the FAZ (red, L3B2) and DNA (blue). Bar = 2 μm. **(I)** TbKin2a-silenced cell at 72 hpi stained for DNA (blue, DAPI) and for nucleoli (green, L1C6). Bar = 2 μm. **(J)** TbKin2a2b-silenced cell at 72 hpi stained for DNA (blue, DAPI) and basal bodies (green, BBA4). Bars = 2 μm. **(F-J)** All images are of cells that are F/M-fixed (except panel F which are M-fixed). Images in (F, J) were taken using DIC microscopy, and/or epifluorescence microscopy. (G, H, I) are 2D projections from a 3D deconvolved stack. Arrowheads: red = bundled flagella phenotype (failed cytokinesis initiation); green = basal body-related abnormalities. Symbols: k^†^ = abnormal kinetoplast (including partially separated kinetoplasts); N^†^ = abnormal nucleus (including unseparated nuclei).

Interestingly, TbKin2a-silenced cells proliferated at a rate 50-70% lower than in uninduced cells after 24 hpi and ceased proliferation by 48 hpi (Fig. 5C). For TbKin2b-silenced cells, proliferation did not decrease significantly for 72-96 hpi (Fig. 5C). Silencing both TbKin2a and TbKin2b together resulted in a cessation of proliferation with kinetics similar to single TbKin2a knockdown (Fig. 5C). Thus, cell proliferation in BSF *T. brucei* requires TbKin2a, but proliferation is not affected by RNAi silencing of TbKin2b.

Evidence of the essential role of TbKin2a in cell proliferation led us to assess its role in cell-cycle progression. We first counted the numbers and observed the morphology of nuclei and kinetoplasts at various time points after RNAi induction (Fig. 5D). Beginning at 24 hpi, the proportion of normal 1N1K (1 nucleus, 1 kinetoplast), 1N2K, and 2N2K cells decreased progressively, while abnormal cells with multiple nuclei and kinetoplasts (yNxK) or with one abnormally large nucleus and multiple kinetoplasts (1†NxK), increased progressively (Fig. 5D). We next observed the effect of TbKin2a silencing on the cell-cycle state by FACS (Fig. 5E). From 24 hpi to 48 hpi, the percentage of cells with 2C DNA content decreased, whereas the percentage of cells with 4C DNA content increased. After 48 hpi, there was a significant increase in the proportion of cells with **≥** 4C DNA content. This suggests that nuclear DNA synthesis continued without complete nuclear separation and/or cytokinesis.

At the morphological level, cells induced for 24 h or longer became much larger than uninduced cells, accumulating multiple nuclei, flagella (Fig. 5F, G), and FAZ (Fig. 5H). Typically, flagella and FAZ clustered at the anterior ends of undivided cells, and there was little evidence of cleavage furrow formation or progression (Fig 5G, H). These results suggest that TbKin2a is important for early cytokinesis, which initiates at the anterior end (Sherwin and Gull, 1989).

In addition, at 24 hpi, 48 hpi, and 72 hpi, cells showed nuclei that were often poorly defined or separated, resulting in large undivided or partially divided nuclear masses with numerous nucleoli (Fig. 5G-J). Although kinetoplast DNA replication appeared to continue, approximately 10-15% of kinetoplasts appeared to be partially separated or not separated (Fig. 5G-J). Basal body duplication and separation were also ongoing, but we observed multiple TbKin2a- and TbKin2a/2b-silenced cells with unseparated kinetoplast DNA or basal bodies having no associated kinetoplast DNA (Fig. 5I-J). The ratio of basal bodies/flagella remained approximately 1:1. These results suggest that TbKin2a contributes to separation of nuclei and/or kinetoplasts and may play a role in the association of basal bodies with kinetoplasts.

### TbKin2a and TbKin2b act additively in forming full-length flagella, but only TbKin2a is required for motility

Considering the established role of kinesin-2 in flagellar assembly in other organisms, we evaluated the effect of silencing TbKin2a and/or TbKin2b on flagella length. Surprisingly, given their sequence divergence from other kinesin-2 proteins, silencing either TbKin2a or TbKin2b for 72 h caused a decrease in flagellum length of 16% and 21%, respectively (Fig. 6A). Silencing both together caused a 42% decrease in flagellar length, which is approximately the additive impact of the individual knockdown. We did not observe a sub-population of flagella that remained long, in contrast with recent observations for silencing IFT proteins in PCF *T. brucei* (Fort et al., 2016) (the cytokinesis defect caused by TbKin2a silencing precludes separate identification of old and new flagella). This indicates that TbKin2a and TbKin2b act additively to build and/or maintain the flagellum in BSF *T. brucei.*

**Fig. 6:**
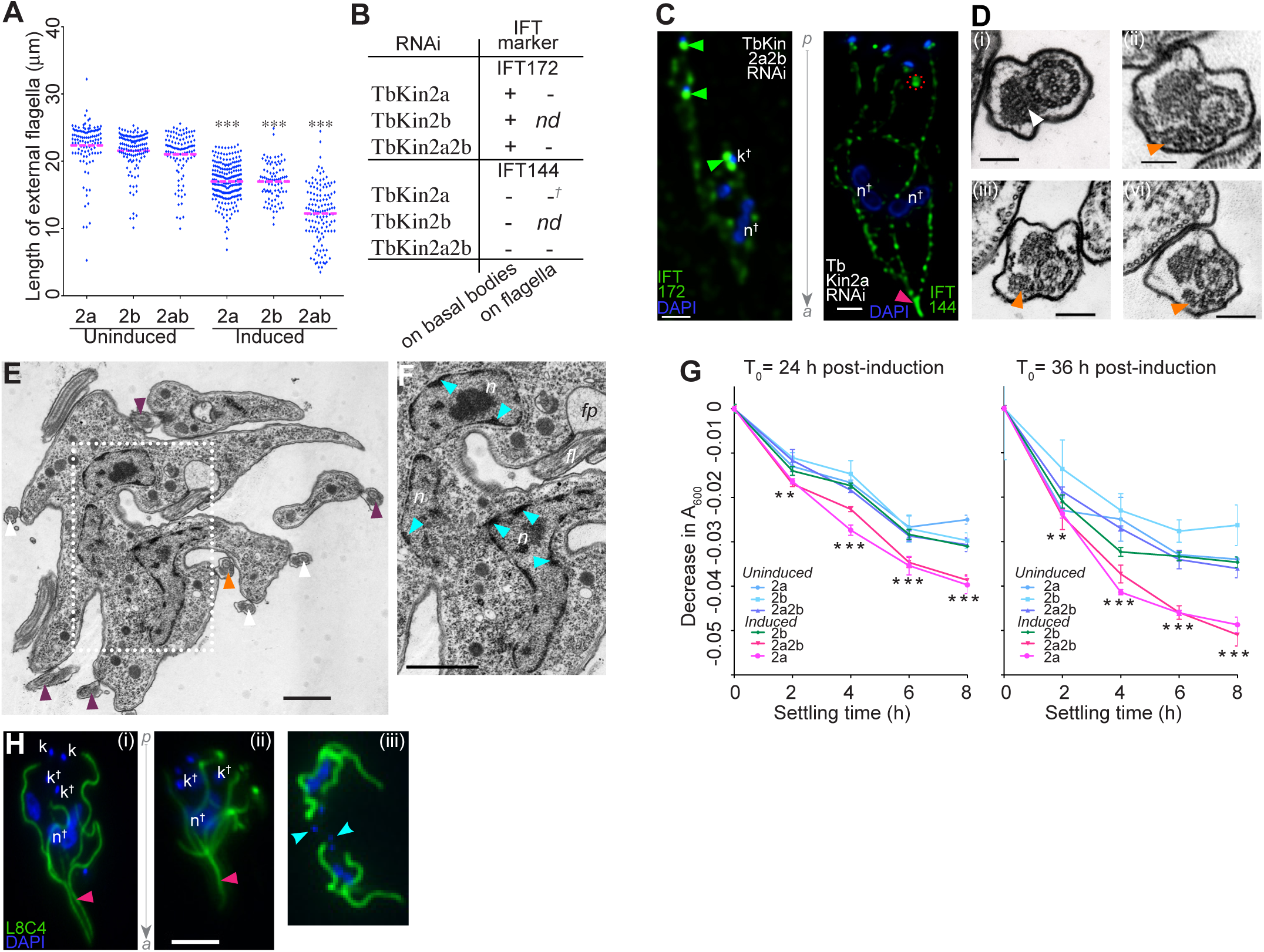
Role of TbKin2a and TbKin2b in flagellar length and motility. **(A)** Flagellar length in uninduced cells (TbKin2a, n = 110; TbKin2b, n = 126; TbKin2a2b, n = 104) and RNAi-induced cells (TbKin2a, n = 211; TbKin2b, n = 87; TbKin2a2b, n = 127) at 48 hpi. Mean = center line, box = SD, whiskers = 95% confidence intervals. 1-way ANOVA with Bonferroni’s multiple comparison tests, ^∗∗∗^ indicates p < 0.001. **(B)** Table indicating whether IFT172 or IFT144 had typical (+) or diminished (-) staining near basal bodies or on flagella in cells induced for RNAi of TbKin2a, TbKin2b, or TbKin2a/2b at 48 hpi. **(C)** 2D projection from a 3D deconvolved stack showing examples of RNAi-impacted IFT protein staining noted in (B) for (left) TbKin2a2b RNAi of IFT172 (compare with uninduced in Fig. 3(D)), and (right) TbKin2a RNAi of IFT144 (compare with uninduced Fig. 3(E)). Gray arrow between images indicates direction of cell anterior/posterior ends. **(D)** Transmission electron micrograph (TEM) of flagellar axonemes for wild type (i) and TbKin2a RNAi cells 48-72 hpi (ii-iv). Bars = 100 nm. (E) TEM of a TbKin2a RNAi cell at 48-72 hpi. **(F)** Magnified view of boxed area from **(E)**. Cyan arrowheads = persistent DNA plaques at inner periphery of nuclear envelope. Symbols: n = nucleus, fp = flagellar pocket and fl = flagellum. Bars = 500 nm. **(G)** Sedimentation assays initiated at 24 and 36 hpi in which the difference in A_600nm_ between matched samples of freshly agitated and settled cells is plotted versus settling time. Data is from 2 independent biological experiments with two technical replicates per experiment. Error bars = SD. 2-way ANOVA with Bonnferroni’s post tests, ^∗∗^ p < 0.01 and ^∗∗∗^ p < 0.001. **(H)** Kinetics of abnormal cell phenotype emergence during early time points (16–18 hpi). Epifluorescence images showing early abnormal phenotypes for TbKin2a RNAi cells stained for the PFR (green, L8C4) and DNA (blue, DAPI) at 16–18 hpi, prior to emergence of abnormal motility in sedimentation assay (G). (i) At 16 hpi and (ii) at 18 hpi show cells with bundled flagella (failed cytokinesis initiation) phenotype. (iii) at 18 hpi shows cells with late cytokinesis (scission) failure. Far posterior ends (cyan arrowheads) of daughter cells are still connected; however, both cells have progressed to the 2K1N^∗^ stage of the next cell cycle. Gray arrow indicates direction of cell anterior and posterior ends. Bar = 5 μm. **(C-F, H)** Arrowheads: red = bundled flagella phenotype (failed cytokinesis initiation); green = basal body abnormalities; orange = abnormal IFT material accumulated; white = normal (no abnormalities); dark red = status could not be determined.

We also evaluated the effect of silencing TbKin2a, TbKin2b or both on the localization of IFT172 and IFT144 to flagella and the region of the basal body (Fig. 6B, C). The staining of IFT172 and IFT144 in flagella was diminished upon silencing TbKin2a or TbKin2a2b. Moreover, the localization of IFT144 near the basal body was diminished upon silencing TbKin2a or TbKin2b, whereas IFT172 staining in this location was unaffected by silencing of either protein. Together with the colocalization of TbKin2a with IFT proteins, these data suggest that TbKin2a, and possibly TbKin2b, play a role in IFT.

To determine whether the defects in flagellar length and IFT protein localization caused by TbKin2a silencing were linked with gross ultrastructural defects in the flagellum, we examined TbKin2a RNAi cells at 48-72 hpi using transmission electron microscopy (TEM) (Fig. 6D). Although cell morphology was dramatically perturbed, we observed infrequent gross structural anomalies in flagella and no FAZ defects. In TbKin2a-silenced cells, 13% (n = 55 images) had abnormal accumulations of material between the flagellar membrane and axoneme outer pairs 3-4 or 8-9 (the location of IFT cargo trains in *T. brucei* (Absalon et al., 2008)), whereas <2% (n = 93 images) of control cells had such accumulations. Moreover, the nuclei of TbKin2a-silenced cells frequently displayed irregular shapes, areas of NE disorganization, and electron dense plaques of various sizes on the inner periphery of the NE (Fig. 6E, F). The plaques were similar in number and location to chromatin plaques observed at the periphery of nuclei in *T. brucei rhodesiense* beginning in late mitosis and persisting into interphase (Farr and Gull, 2012; Vickerman and Preston, 1970). This ultrastructural evidence is consistent with the conclusion that TbKin2a plays a role in IFT and suggests that it may also play a role in late cell-cycle timing as well as NE and chromatin organization.

To measure the impact of TbKin2a and TbKin2b silencing on flagellar motility, we adapted a previously established sedimentation assay for PCF *T. brucei* in which the rate of sedimentation is inversely proportional to the motility capacity (Bastin et al., 1999; Ralston et al., 2006). Sedimentation behavior was assessed over 4-8 h, beginning at 18 hpi, 24 hpi and 36 hpi. At 18-22 hpi, sedimentation rates for induced and uninduced cells were statistically equivalent (not shown). However, after 24 hpi, cells silenced for TbKin2a or TbKin2a/2b sedimented significantly faster than uninduced controls at every time point considered (Fig. 6G). In contrast, silencing TbKin2b alone did not result in faster sedimentation relative to controls. We conclude that TbKin2a silencing, but not TbKin2b silencing, results in a significant, progressive decrease in motility.

Finally, we evaluated the kinetics of onset of the TbKin2a RNAi phenotype by identifying abnormal cell morphologies as they formed over the first 26 hpi. We observed small numbers of abnormal cells beginning 16 hpi, and over the next 4-8 hpi abnormal cells accumulated such that by 26 hpi over 20% of the cell population had abnormal morphologies. The most frequent abnormal phenotype was bundled flagella at the anterior end (Fig. 6H(i) and (ii)), and we also observed cells that were attached at their far posterior ends (Fig. 6H(iii)). Thus, two of the earliest phenotypes associated with silencing TbKin2a are an apparent failure to initiate or failure to complete cytokinesis.

## Discussion

Kinesin-2 proteins perform an important and conserved function in IFT and flagellar biogenesis of primary and motile cilia (Lechtreck, 2015; Scholey, 2013). Kinesin-2 proteins have also been implicated in bidirectional endoplasmic reticulum (ER) to Golgi transport (Brown et al., 2014; Stauber et al., 2006), endosomal trafficking (Granger et al., 2014), transport in neurons (Hirokawa et al., 2010), cilium-based signaling (Goetz and Anderson, 2010), chromosome segregation (Haraguchi et al., 2006; Miller et al., 2005), and cytokinesis completion (Brown et al., 1999; Fan and Beck, 2004). Here, we report that the two kinesin-2 proteins TbKin2a and TbKin2b in BSF *T. brucei* function in flagellar biogenesis, but only TbKin2a appears to be crucial for cell proliferation. In addition, TbKin2a is found on the FAZ, suggesting a possible role in FAZ-based transport. Thus, kinesin-2 proteins in trypanosomes are likely to perform both canonical and trypanosome-specific roles.

### Trypanosome kinesin-2 proteins have divergent NST sequences

The results of our MEME analysis complements previous kinesin bioinformatics studies (Berriman et al., 2005; Wickstead et al., 2010b) by identifying conserved sequence motifs in the non-motor-domain NST portions of kinesin-2 proteins. Unexpectedly, we found that kinesin-2 proteins contain over 20 conserved NST motifs that can be assigned into several key motif groups and are broadly shared across all superfamilies of flagellated eukaryotes. Several of these motifs had been previously recognized within a narrower phylogenetic context (De Marco et al., 2001; De Marco et al., 2003; Doodhi et al., 2009; Imanishi et al., 2006; Vukajlovic et al., 2011), while others, in particular NST motifs specific to subgroups 2A, 2B, and 2C, appear not to have been observed previously. Some motifs and motif groups were common to almost all kinesin-2 taxa, while others were specific for heterotrimeric kinesins 2A/2B or homodimeric kinesin-2C taxa.

Although the IFT machinery is highly conserved within the kinetoplastids (van Dam et al., 2013), it was surprising that the kinetoplastid taxa did not share common kinesin-2 NST motif groups and had little evidence of conserved individual motifs (Fig. S2). Kinetoplastids are also the only kinesin-2 containing organisms identified that do not encode non-motor KAP homologs in their genomes (Julkowska and Bastin, 2009). The free-living kinetoplastid *B. saltans* also does not encode a KAP homolog, suggesting that it was not lost as a result of parasitism. Our MEME analysis and KAP genomic data do not support bioinformatic assignment of kinetoplastid kinesin-2 taxa as orthologs of the canonical heterotrimeric kinesin-2A or 2B forms. However, a kinetoplastid kinesin-2 protein did share a single significant kinesin-2C motif with Metazoa, which was the sole example of a homodimeric-2C motif among non-holozoan single-celled organisms. Together with the markedly different phenotypes caused by TbKin2a versus TbKin2b RNAi silencing, these data suggest that kinesin-2 proteins in *T. brucei* do not form canonical heterotrimers, but instead represent a distinct subgroup within the kinesin-2 family, and form homodimers or heterodimers that differ significantly in sequences and motifs and may differ in function compared with kinesin-2 proteins in other organisms.

### Trypanosome kinesin-2 proteins are important for flagellum biosynthesis

We found that silencing TbKin2a or TbKin2b results in a ~ 20% decrease in flagellum length, while silencing both causes a decrease of 40% (although flagella are present). Moreover, TbKin2a localizes along the flagellum, with a concentration on basal bodies and the proximal flagellum, similar to the localization of *C. reinhardtii* kinesin-2 FLA10 (Cole et al., 1998; Deane et al., 2001; Vashishtha et al., 1996). TbKin2a also colocalizes with both IFTB protein IFT172 and IFTA protein IFT144 near basal bodies. Silencing TbKin2a decreased staining of both IFT172 and IFT144 on the flagellum and decreased the localization of IFT144, but not IFT172, to the basal body. Moreover, flagella in TbKin2a-silenced cells showed accumulation of electron-dense material along axonemes, but otherwise did not show gross flagellar abnormalities (we could not always resolve fine structural elements such as inner dynein arms, which require kinesin-2 for transport in *C. reinhardtii* (Piperno and Mead, 1997; Piperno et al., 1996)). These data suggest that TbKin2a and TbKin2B participate in IFT.

One distinction between TbKin2a and TbKin2b that is of potential significance is the presence of a putative C-terminal nuclear localization signal sequence in TbKin2a, but not in TbKin2b (NLS; predicted using NucPred (Brameier et al., 2007) and MultiLoc2 (Blum et al., 2009)); this sequence might also function as a ciliary localization signal sequence (CLS) (Verhey et al., 2011). For the homodimeric kinesin-2 protein KIF17, an NLS/CLS mediates entry into the ciliary compartment (Dishinger et al., 2010; Kee et al., 2012). Thus, TbKin2a and TbKin2b may play different roles in cargo entry into flagella. Recent investigations in *C. elegans* indicate that heterotrimeric kinesin-2A/2B is primarily responsible for cargo entry into the flagellum transition zone. Following cargo entry, kinesin-2A/2B gradually undocks while homodimeric kinesin-2C docks in the proximal segment. Finally, kinesin-2C becomes the primary transport motor in the distal segment (Prevo et al., 2015). By analogy, TbKin2a and TbKin2b may play different roles in cargo transport at different regions of the flagellum.

In many other organisms, kinesin-2 proteins are essential for both flagellum biogenesis and maintenance (Lechtreck, 2015; Scholey, 2013). However, in BSF *T. brucei* silencing kinesin-2 proteins resulted in partial but not complete inhibition of flagellum biosynthesis and maintenance. This may be due to incomplete RNAi silencing, slow protein turnover, the presence of other kinesins that participate in flagellum biosynthesis and maintenance, or increasingly pleiotropic effects of RNAi over time on other processes in BSF cells such as cell-cycle progression, which result in inhibition of cellular functions, or cell death before flagellar shortening is complete (Blaineau et al., 2007; Broadhead et al., 2006; Chan and Ersfeld, 2010; Demonchy et al., 2009; Marande and Kohl, 2011; Wickstead et al., 2010a). Interestingly, in PCF *T. brucei* a recent study revealed that IFT is active in both new and old flagella, but that silencing either anterograde IFTB or retrograde IFTA components caused a failure in new flagellum biosynthesis (although with somewhat different phenotypes) but no defect in old flagellum maintenance (Fort et al., 2016). Our results are consistent with a role for kinesin-2 proteins in anterograde IFT during new flagellum biosynthesis. However, the fact that we do not observe a sub-population of flagella that remain long in TbKin2a/TbKin2b-silenced BSF *T. brucei* suggests that TbKin2a and TbKin2b may be important for flagellar maintenance in BSF cells.

### TbKin2a is essential for cell proliferation

Interestingly, silencing TbKin2a expression caused a cessation of cell proliferation beginning at ~ 24 hpi, whereas silencing TbKin2b expression had no apparent effect on cell proliferation. Silencing TbKin2a also caused a failure in cytokinesis, with the accumulation of greatly enlarged cells with multiple nuclei, kinetoplasts, basal bodies, FAZ, and flagella, as well as decreased cell motility. It is surprising that only TbKin2a is important for proliferation, cytokinesis and motility, although both TbKin2a and TbKin2b contribute to flagellar length. This suggests that flagellar length is flexible to decreases of at least 20% and that shorter flagella in BSF cells can be functional, perhaps through a compensatory mechanism such as decreased cell size, as in PCF *T. brucei* (Kohl et al., 2003).

The cell proliferation defects caused by silencing TbKin2a in BSF cells are similar at a gross level to the defects caused by RNAi silencing of other flagellar proteins, FAZ proteins, and basal body proteins (Broadhead et al., 2006; LaCount et al., 2002; Morris et al., 2001; Ralston et al., 2006), as well as cell-cycle-related proteins, other signaling proteins, and cell-surface proteins (reviewed in (Farr and Gull, 2012; Hammarton et al., 2007b)). The commonality of phenotypes is specific to BSF cells and may be due to the fact that silencing these factors directly or indirectly impacts cytokinesis (Hammarton et al., 2007b; Zhou et al., 2014). It remains possible that, in PCF cells, the functions of TbKin2a and TbKin2b are distinct.

It has been hypothesized that cell proliferation and cytokinesis defects in BSF trypanosomes can result from inhibited flagellar beating (Broadhead et al., 2006; Ralston and Hill, 2006). However, defects in flagellar beating may not explain the cell proliferation and cytokinesis phenotypes observed in TbKin2a-silenced cells for several reasons. First, normal flagellar beating is not essential for cytokinesis, because the cell proliferation and cytokinesis defects caused by silencing dynein light chain 1 by RNAi can be rescued by the expression of site-specific mutants that are still defective in flagellar motility (Kisalu et al., 2014; Ralston et al., 2011). Second, silencing TbKin2a did not cause gross ultrastructural abnormalities in flagella or the flagellar pocket other than abnormal IFT train accumulation, in contrast to the major defects caused by silencing other flagellar proteins PFR2, TAX-1, TAX-2, DIGIT, or TbMBO2 (Broadhead et al., 2006; Farr and Gull, 2009). Third, we find that following TbKin2a RNAi induction, the kinetics of onset of the cytokinesis defect precedes the defect in cell motility. For example, at 16-20 hpi, abnormal cells with bundled flagella and FAZ at their anterior ends are observed, suggesting a defect in cytokinesis initiation. Abnormal cells are also observed connected at their posterior ends, suggesting a defect in cell scission. At 24 hpi, cell proliferation is affected, and there is a decrease in motility although the flagella continue to beat. These considerations suggest that TbKin2a may play a role in cytokinesis initiation that is independent of any role in flagellar motility. This may be due to its involvement in transporting cargoes into flagella that themselves are important for cytokinesis or its on the FAZ, as discussed below.

### Implications of TbKin2a localization on the FAZ

It is intriguing that TbKin2a also localizes to the FAZ. However, TbKin2a does not appear to be important in the construction, maintenance, or structure of the FAZ because we did not observe abnormalities in the FAZ or flagellar detachment in TbKin2a or TbKin2a2b-silenced cells. Instead, we propose that TbKin2a interacts with the FAZ via the MtQ and may transport other cargoes, including those essential for cell division. This also the simplest model, given the ability of TbKin2a to bind to microtubules. Because MtQ microtubules are uniformly oriented with their plus ends towards the cell anterior (Gull, 1999), antiparallel to the remainder of the subpellicular array (Robinson et al., 1995), they represent a potential track for moving cargoes along the anterior-posterior cell axis.

We speculate that cargoes transported by TbKin2a along the FAZ could include both organelles and cell cycle regulatory molecules. Recent studies have revealed that the FAZ is linked with other single-copy organelles including the kinetoplast, tripartite attachment complex (TAC), basal body, flagellum, flagellar pocket collar, and bilobe structure with ER and Golgi exit sites (Gadelha et al., 2009; Gheiratmand et al., 2013; Gluenz et al., 2011; Lacomble et al., 2009; Lacomble et al., 2010; Sunter et al., 2015; Zhou et al., 2015). TbKin2a may be involved in transporting components involved in these interactions. For example, basal bodies and kinetoplasts are connected by the TAC (Ogbadoyi et al., 2003), and basal body duplication and separation are necessary for kinetoplast segregation (Robinson and Gull, 1991). We observed defects in kinetoplast separation and basal body to kinetoplast attachment in TbKin2a-silenced cells, suggesting that TbKin2a may participate in TAC assembly or maintenance. TbKin2a may also be important for the transport of structural or regulatory proteins that are important for cell cycle regulation as well as duplication and segregation of many of the above-mentioned components, which include the centrins 1, 2, and 4 (Selvapandiyan et al., 2007; Shi et al., 2008; Wang et al., 2012) as well as polo-like kinase (TbPLK) (de Graffenried et al., 2013; de Graffenried et al., 2008; Ikeda and de Graffenried, 2012; Li et al., 2010; Umeyama and Wang, 2008). Notably, TbPLK is transported along the FAZ to the cell’s anterior tip (Ikeda and de Graffenried, 2012; Li et al., 2010; Sun and Wang, 2011; Yu et al., 2012), where it is required for cytokinesis initiation (Hammarton et al., 2007a; Kumar and Wang, 2006). Future experiments will be aimed at determining whether TbKin2a and TbKin2b transport distinct cargos along the flagellum and/or FAZ, and how cargo transport contributes to flagellum construction, organelle transport and positioning, and cytokinesis.

## Materials and Methods

### MEME suite analysis

Multiple EM for Motif Elicitation (MEME) web-based software (Bailey and Elkan, 1994; Bailey and Gribskov, 1998; Bailey et al., 2009; Bailey et al., 2006) (versions 3.91-4.11) was used to analyze the NST domains of putative kinesin-2 proteins. Using kinesin-2 taxa identified in a phylogenetic analysis of kinesin motor domain sequences from 45 diverse organisms (Wickstead et al., 2010b) as starting point, we carried out more than 80 independent MEME computational runs including multiple combinations of kinesin-2 NST data subsets and input parameters with multiple controls. To ensure that all NST domains consistently included a full neck region, which has been shown to evolve independently of the motor domain (Case et al., 2000; Vale and Fletterick, 1997), our NST domain for each taxon began with the final 3 amino acids (positions 343-345) of the PFAM-defined motor domain sequence specification (PFAM00225). MEME results were sensitive to changes in inputs and assumptions including composition, length and number of protein sequences, expected motif distribution and run parameters. To overcome this sensitivity, many runs of kinesin-2 sequences as well as control sequences were conducted until consistent result patterns were observed. Our controls included running a full MEME analysis of: kinesin-1 family NST sequences; kinesin-1, -3 and -4 family neck domain sequences; kinetoplastid and non-kinetoplastid kinesin NST sequences; and scrambled kinesin-2 NST sequences (NST sequences are found in Supplemental Table S4). Overall results were genrally robust and consistent but as already noted, could vary from run to run. Kinesin-2 NST motif results presented here are from a single (MEME 83) that we judged to be highly consistent with overall result patterns. All MEME statistics presented were generated by standard MEME programs (Bailey and Elkan, 1994; see Supplemental Figures S1-S2, Supplemental Table S3).

### Cell culture and RNA interference

BSF *T. brucei brucei* Lister 427-derived cell line 90-13 (Wirtz et al., 1999) was cultured and maintained as described previously (Li and Wang, 2006; Tu and Wang, 2004), with splitting of the cultures at final densities of ~1.0 × 10^6^ cells/ml. For RNAi, DNA corresponding to 380-bp *T. brucei* gene Tb927.5.2090 (TbKin2a) or 462-bp gene Tb927.11.13920 (TbKin2b) (selected using RNAit software; (Redmond et al., 2003)) were amplified by PCR from genomic DNA (primers listed in Table S5). For RNAi of TbKin2a or TbKin2b, PCR fragments were ligated into the XhoI and HindIII sites of pZJM (Wang et al., 2000). For RNAi of both TbKin2a and TbKin2b, the TbKin2b PCR product was ligated into the XbaI and XhoI sites in pZJM that already contained TbKin2a. All DNA sequences were verified. Transfections (Burkard et al., 2007; Li and Wang, 2006) and selection of monoclonal transformants were performed using agarose plates as described previously (Carruthers and Cross, 1992; Tan et al., 2002). For induction of RNAi, monoclonal transformants were cultured with 1 μg/ml tetracycline for 0-96 h as indicated. A list of primers used is found in Supplemental Table S5. For growth curves, live cells were counted using a hemocytometer.

### Antibody generation and immunoblotting

A portion of the TbKin2a gene encoding amino acids 391–696 plus a C-terminal 6XHis tag (TbKin2a.391-696-His) was amplified by PCR using primers listed in Table S5, ligated into the NcoI and NotI sites in pET22b(+) (Novagen, Madison, WI), and verified by DNA sequencing. Recombinant TbKin2a.391-696-His was expressed in *E. coli* BL21-CodonPlus-RP cells (Agilent Technologies, Cedar Creek, TX) after induction with 400-μM IPTG at 37°C for 2 h. The protein was purified by Ni-NTA (Qiagen, Valencia, CA) affinity chromatography followed by gel filtration chromatography on a Superdex 75 column (GE Healthcare, Piscataway, NJ). To generate anti-TbKin2a antibody, rabbits were immunized by Covance Inc. (Princeton, NJ) using purified TbKin2a.391-696-His. Antibody was affinity purified using standard procedures. For immunoblotting, affinity-purified anti-TbKin2a antibody was used at a dilution of 1:10,000, and YL 1/2 rat monoclonal anti-tyrosylated α-tubulin antibody (Millipore, Billerica, MA) and/or KMX mouse anti-ß-tubulin antibody (Birkett et al., 1985) were used at a dilution of 1:100.

### RT-PCR and qRT-PCR

For all RT-PCR and qRT-PCR studies, total RNA (from at least two independent biological replicates) was extracted and isolated individually for each experiment from freshly spun-down, unwashed *T. brucei* BSF RNAi cells (induced or uninduced) using either TRI Reagent or RNAzol (Molecular Research Center (MRC), Cincinnati OH). RNA for semi-quantitative RT-PCR was purified after isolation using MRC RNA precipitation protocols, including a DNAse treatment step using TurboDNAse (Thermo Scientific). For quantitative qRT-PCR from RNAi cells, RNA extracted and isolated as above from n = 4 (TbKin-2a), n = 2 (TbKin2b), and n = 4 (TbKin2a2b) biological experiments was individually purified either as above, or by immobilizing extracted and isolated RNA on silica columns (Direct-zol, Zymo Research (ZR), Carlsberg, CA), treating with DNAse, and purifying, all as described in the manufacturer’s protocols (except that an additional wash step of 100% ethanol was included) and as a final step, eluted with pure DEPC water. Purified RNA in DEPC water was analyzed using a Nanodrop spectrophotometer (Thermo Scientific) and stored at -80°C. For qRT-PCR, purified RNA integrity for each sample was confirmed using an Agilent Bioanalyzer 2100 (Agilent Technologies, Santa Clara, CA) at the UC Berkeley QB3 Functional Genomics Laboratory. All primers were from Integrated DNA Technologies (Coralville, IA) (Table S5). For semi-quantitative RT-PCR, first-strand cDNAs were generated from RNA using SuperScript III First-Strand Synthesis System (Thermo Scientific) with oligo(dT) primers, and PCR was performed using first-strand cDNA and with gene-specific primers. qRT-PCR used a two-step procedure with first-strand cDNAs generated from purified RNA using a Superscript VILO cDNA Synthesis Kit (Thermo Scientific). We evaluated *T. brucei* potential reference standard genes as recommended previously (Brenndörfer and Boshart, 2010). Only one gene, TERT (telomerase reverse transcriptase, Tb927.11.10190), had minimal expression variation under our experimental conditions and was selected as sole reference standard. All qPCR work was performed in 96-well plates using either Applied Biosystems (ABI) 7500 Fast Real-Time PCR System and iTaq SYBR Green Supermix (Thermo Scientific); or Roche LightCycler96 System using iQ SYBR Green Master Mix (BioRad, Inc., Richmond, CA) using separate wells for reference and target reactions. Primers for qRT-PCR were designed using ABI Primer Express Version 2.0 software and/ or Primer3 software (Untergasser et al., 2012). Amplification efficiency of each gene/primer combination was confirmed over a 4- to 5-log dilution series. Relative gene expression for qRT-PCR was determined for each biological experiment, and results were aggregated using the 2^−ΔΔC^T method (Schmittgen and Livak, 2008).

### Fluorescence activated cell sorting

Cell samples for fluorescence-activated cell sorting (FACS) were prepared as described previously (Tu and Wang, 2004). FACS was carried out using an Epics XL flow cytometer (Beckman Coulter, Brea, CA). Data were analyzed using FlowJo 4-5 software (Tree Star, Ashland, OR).

### Motility assays

At experiment initiation, cells were centrifuged and transferred to 1–10 ml of fresh settling media (a 50/50 v/v mix of heat-treated tetracycline-free fetal bovine serum (Atlanta Biologicals, Flowery Branch, GA) and regular complete HMI-9 media (Engstler et al., 2007) at starting densities of ~ 0.5–3 × 10^5^ cells/ml, with or without tetracycline at 37°C with 5% CO_2_. Using previously described methods, including media replacement at 12-24 h intervals (Hesse et al., 1995), un-induced cells were grown in settling media for up to 48 h and reached final densities of up to ~ 3 × 10^7^ cells/ml. At 18, 24 or 36 h after growth in the settling media and RNAi induction, cells were counted and 1 ml of cells was transferred into each of 2 paired cuvettes and sealed with sterile, gas-permeable film. Cuvettes were agitated and the initial A_600nm_ at 0 min was measured using a SpectraMax M2 multi-detection reader (Molecular Devices, Sunnyvale, CA) or Genesys 10 spectrophotometer (Thermo Fisher Scientific, Waltham, MA). Cuvettes were incubated at 37°C with 5% CO_2_, and at each subsequent time point (every 1–2 h up to 8 h), cuvettes were removed from the incubator, and one of each pair was agitated again, while the other remained still. A_600nm_ measurements were obtained and differences in absorbance were calculated.

### Staining for fluorescence microscopy

Except for counts of nuclei/kinetoplast morphology, for all immunofluorescence microscopy, live cells were either (i) centrifuged at 600-1200 x g and diluted in 37°C fresh HMI-9 complete media or HBS/G (25 mM HEPES pH 7.1, 140 mM NaCl, 5 mM KCl, 0.75 mM Na_2_HPO_4_, 22 mM glucose) to 1.5–2.0 × 10^6^ cells/ml, or (ii) if too fragile to centrifuge (for some RNAi cells), then cells at 0.5–1.2 × 10^6^ cells/ml were settled directly onto poly-L-lysine (Amanda Polymers, Birmingham, AL) coated coverslips (#1.5 high precision-type). For F/M-type fixation, cells on coverslips were washed 1–2× in HBS/G, then fixed immediately in HBS/G buffer with freshly added formaldehyde (Ted Pella, Inc.) at 0.75% to 4% for 5–10 min at room temperature, washed 2–3x in HBS/G, placed on ice, and then fixed with -20°C methanol for 5–7 min, followed by 3–5× washes in PEME (100 mM PIPES pH6.9, 2 mM EGTA, 1 mM Mg_2_SO_4_, 0.5 mM EDTA) (Robinson et al., 1991) and a 5–10 min rehydration in PEME. For methanol-only fixation (including flagellar length or intensity measurements), cells on coverslips were washed 1–2× in HBS/G, placed on ice and fixed in -20°C 100% methanol for 10–20 min, followed by PEME washing and rehydration steps as above. Detergent extraction was carried out before fixation on live unfixed cells at room temperature using the Nonidet P-40 method described previously (Robinson et al., 1991), except that live cells were settled onto and extracted on poly-L-lysine–coated coverslips; IGEPAL CA-630 was substituted for Nonidet P-40 at equal concentrations; extracted cytoskeletons were stabilized in 2× cOmplete™ protease inhibitor cocktail, EDTA-free (Roche Diagnostics), plus 1 mM E 64d cysteine protease inhibitor (Apex Biotechnology, Taiwan). All buffers used prior to fixation were supplemented with 5 mM ATP (molecular biology grade, Sigma) and 2 mM MgCl_2_. Cells were permeabilized, blocked and incubated in primary and secondary antibodies as described (Sagolla et al., 2006) at the following concentrations: L8C4 (Kohl et al., 1999), 1:100; KMX (Birkett et al., 1985), 1:25-1:50; L3B2 mouse anti-FAZ (Kohl et al., 1999), 1:50; YL1/2 (EMD Millipore, Darmstadt, Germany), 1:100; BBA4 mouse monoclonal anti-basal-body (Woods et al., 1989), 1:50; L1C6 mouse anti-nucleolar protein (Durand-Dubief and Bastin, 2003), 1:200; mouse anti-IFT144 (PIFTF6) (Absalon et al., 2008), 1:200; mouse anti-IFT172 (Absalon et al., 2008), 1:1,000; rabbit anti-TbKin2a (this study), 1:20,000. Secondary antibodies conjugated to AlexaFluor 488 or AlexaFluor 568 (Invitrogen) were diluted at 1:400 with DAPI at 2.5 ng/ml. For double staining with mouse anti-IFT172 and mouse L3B2, anti-IFT172 was labeled with Zenon AlexaFluor 488 (Invitrogen) and used at a 1:1000 dilution, and L3B2 was labeled using Zenon AlexaFluor 568 (Invitrogen) and used at 1:10, using the recommended Zenon protocol. Coverslips were mounted onto slides using ProLong Gold antifade.

For nuclei and kinetoplast morphology counts, 0.7–2.0 × 10^7^ live cells were centrifuged, washed in PBS/G (PBS with 20 mM glucose) then fixed with 4% formaldehyde for 20–30 min. Fixed cells were isolated by centrifugation, washed in PBS, placed on poly-L-lysine-coated coverslips and post-fixed with -20°C methanol for 20–30 min. Fixed cells on coverslips were permeabilized and blocked in PBS with 1% bovine serum albumin (BSA) and 0.1% Triton X-100 for 1 h. For staining, cells were incubated with mouse anti-PFR L8C4 antibody (Kohl et al., 1999) at 1:50 in PBS with 1% BSA and then with AlexaFluor 488 anti-mouse secondary antibody (Invitrogen) and DAPI (2.5 ng/ml). Samples were mounted using ProLong Gold antifade (Invitrogen) and allowed to hard set in dark at 4° C for at least 24 h, and sealed with clear nail polish before imaging.

### Shear Method for separating flagella from FAZ

We devised a simple parallel-plate flow chamber device and shearing procedure to apply measured, consistent shear forces. 24 × 40 mm poly-L-lysine coated #1.5 high-precision coverslips were pre-weighed dry, then fully fixed cells (see staining for fluorescence microscopy) were added, coverslips were mounted onto a cleaned glass slide with 50–120 μl ProLong Gold antifade (Invitrogen), and the assembly was placed into a mounting slot in the device base, which had been levelled and clamped in place and maintained at room temperature. Coverslips were re-weighed after each step above. A 25 x 50 mm glass slide top-plate, with or without attached weights (total top plate mass as used ranged from 7.55g – 10.52g) was loaded carefully onto the coverslip, flush along the back side, between device guides. All masses were determined using a single, calibrated, fully-enclosed Mettler Toledo AB54 S analytical balance. Using a stopwatch and pusher rod placed directly at the back of the coverslip, the coverslip and top plate assembly was pushed by hand in a single direction at a constant velocity of ~0.7 - 1 mm/s for 4-5 s over a measured distance of ~4-5 mm. Immediately before and after the above-mentioned steps, clean, dry pre-weighed Kimwipes (Kimberly-Clark, Roswell, GA) held in micro-forceps were used to blot-up excess liquid and immediately reweighed, with all before and after wipe masses recorded. Shear-treated coverslips were then allowed to set in dark at 4° C, and sealed as per Staining for Fluorescence Microscopy.

### Microscopy and image analysis

Epifluorescence microscopy was performed using an Olympus IX71 microscope with 60× (1.40 NA) and 100× (1.35 NA) PlanApo objectives and a Coolsnap HQ camera (Photometrics, Tucson, AZ). Images were captured using Metamorph (Molecular Devices) or μManager software (Edelstein et al., 2010). Brightness/contrast levels were adjusted using ImageJ and Image-J2/Fiji (Schneider et al., 2012) and Photoshop CS3 software (Adobe, San Jose, CA). For flagellar intensity measurements, ImageJ/J2 were used to measure mean fluorescence intensity at 0.1-mm intervals along 53 individual mature flagella (i.e., data do not include new flagella) on methanol-fixed cells from 3 independent experiments. For flagellar length measurements, ImageJ/J2 were used to measure lengths of PFR (stained with L8C4) in uninduced and induced methanol-fixed cells from 4 independent experiments.

Deconvolution microscopy was carried out using a DeltaVision Elite microscope (GE Healthcare Life Sciences, Pittsburgh, PA) with Olympus UPlanApo 100× (1.35 NA) or UPlanApo 60× (1.40 NA) objectives and Coolsnap ES^2^ camera (Photometrics). Images were captured using SoftWoRx v6 software (GE Healthcare). From 40-200 serial z-sections per channel were acquired at 0.10–0.40-μm intervals. Image z-stacks were deconvolved using Huygens Professional v14 (Scientific Volume Imaging B.V., Hilversum, Netherlands) or SoftWoRx software (“deconvolved stacks”). Images were analyzed using Imaris v7 (Bitplane, Zurich, Switzerland), and Image-J/J2. Two-dimensional projection images were generated from three-dimensional deconvolved stacks using Imaris, ImageJ/J2, Huygens Professional or SoftWoRx. Image adjustments required for presentation were made using ImageJ/J2 and Photoshop. For colocalization analysis with BBA4, IFT172, IFT144, TbKin2a and DNA, Imaris v7 ImarisColoc was used on deconvolved z-stack data, and statistics were computed using Imaris v7 and Prism 6. Threshold intensity levels for each channel were analyzed for relative sensitivity of colocalization statistics to lower boundary thresholds ranging from 2–-20% of maximum intensity, and were generally set at 10% of the maximum intensity for that channel.

For transmission electron microscopy (TEM), cells were processed as previously described (Tu et al., 2005), and were imaged using a JEOL 1200 transmission electron microscope.

### Statistical analyses

Quantitative data was calculated and graphed using Prism 6 (GraphPad Software, La Jolla, CA). Sample sizes (n) were sufficient to detect statistical significance at the levels indicated. For Figure 6A, the relative length statistics were computed using one-way analysis of variance (ANOVA) with Bonferroni’s multiple comparison test in Prism 6. For Figure 6G, settling data statistics were computed using two-way ANOVA with Bonferroni’s post tests in Prism 6. Statistical significance was measured used two-tailed distribution tests.

## Acknowledgements

We particularly thank C.C. Wang, and former members of the Wang lab at UCSF: Ziyin Li, Xiaoming Tu, Stephane Gourguechon, Praveen Kumar, and others, for reagents, procedures, unpublished observations, draft manuscript review (C.C.W. and Z.L.), insights, advice, and encouragement. We thank Philippe Bastin, Benjamin Morga, and Thierry Blisnick of the Pasteur Institute for antibodies to IFT172 and IFT144, unpublished images, insights, and suggestions. We thank Scott Dawson at UC Davis for extensive insights and advice on phylogenetic methods and interpretation. We thank Ryan Case for insights and suggestions regarding bioinformatic analysis of kinesins. We thank the following individuals for reagents: Keith Gull at Oxford University for antibodies (KMX, L8C4, L1B6, L3B2 and BBA4), Paul Englund at Johns Hopkins University for the pZJM vector, and George Cross at Rockefeller University for the 90-13 cell line. We thank Denise Schichnes and Steve Ruzin of the UC Berkeley Biological Imaging Facility for assistance with microscopy, image processing, and colocalization. Finally, we thank Anna Albisetti for comments on the manuscript, and current and former Welch Lab members for their suggestions, support, and patience.

## Author contributions

R.L.D., B.M.H., and M.D.W. conceived the experiments. R.L.D., B.M.H., H.W., R.L.J., J.M. W.Z.C., and M.D.W. designed the experiments. R.L.D., B.M.H., H.W., R.L.J., and J.M. executed the experiments. R.L.D., B.M.H., W.Z.C., H.W., and M.D.W. interpreted the findings being published. R.L.D. and M.D.W. drafted and revised the article.

---

**Fig. S1.**
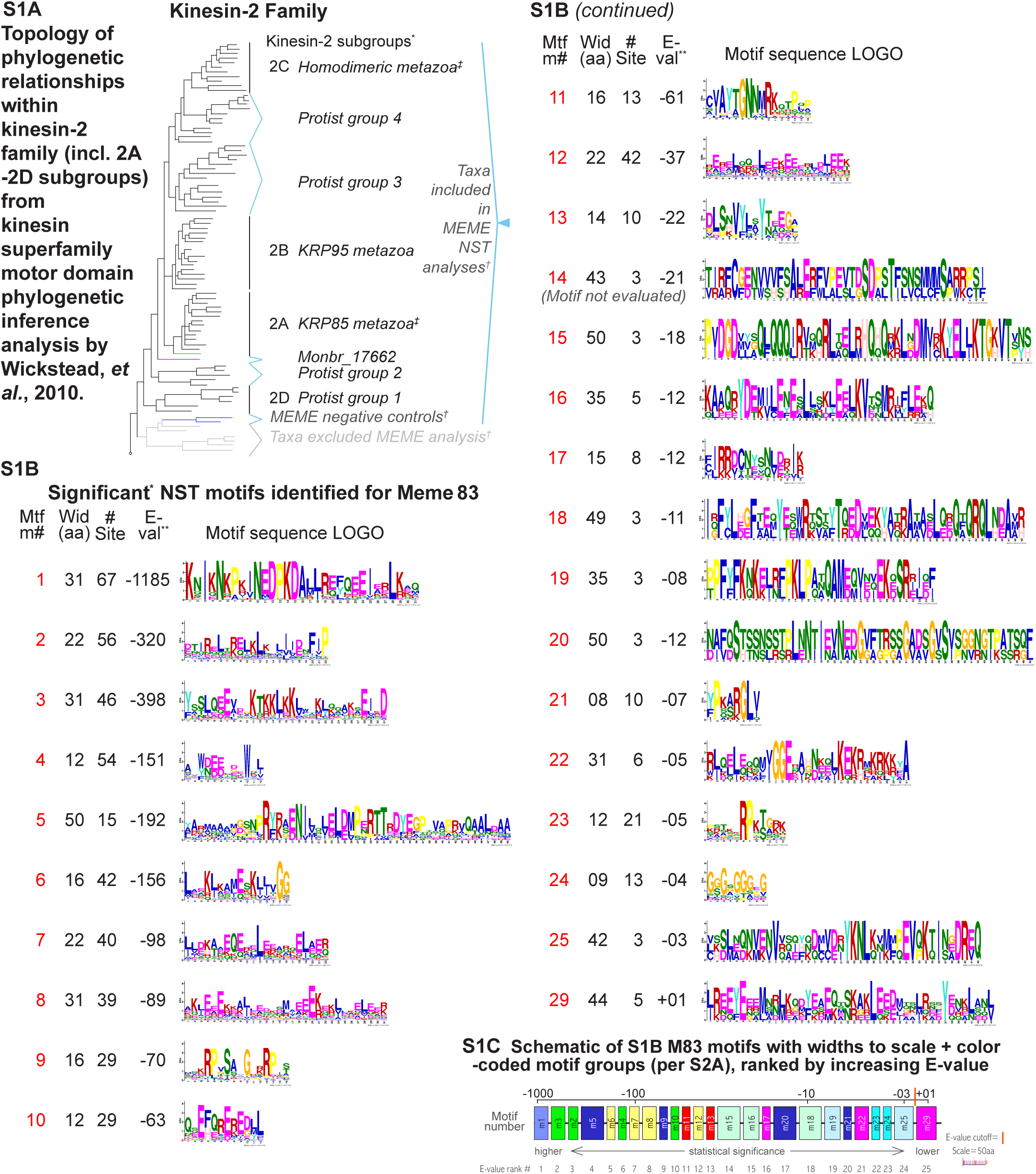
Kinesin-2 family phylogeny and NST motifs. **(A)** Kinesin-2 family phylogentic tree topology adapted from the previous analysis by using motor domain sequences; branch lengths shown do not indicate evolutionary distances. Symbols: ^*†*^Our early MEME analyses showed these NST sequences to be distinct from remaining kinesin-2 taxa; for additional information see Table S3A. ^*‡*^Monbr_16629 and Monbr_23354 of the holozoan *M. brevicolis* were respectively assigned to subgroups 2A and 2C by. Monbr_23354 had a putative NST domain too short for MEME analysis. **(B)** Kinesin-2 motor domain sequences are more highly conserved that kinesin-2 NST sequences (see Tables S1 for motor domain sequence comparisons, and Tables S2 for NST sequence comparisons). The NST analyses relied primarily upon the MEME tool suite, which can identify conserved motifs independently of sequence alignment or motif order. We carried out over 80 MEME runs because of variation between runs due to differences in run parameters and taxa used as described in Materials and Methods. Shown in the figure are the 25 statistically significant NST motifs (based on MEME-determined individual motif overall E-values) plus one additional motif identified in a single representative run (MEME 83). Shown are: motif number (Mft m#); optimal width (amino acids or aa); the total number of significant, non-overlapping, independent motifs identified in the data set (# sites); the E-value overall statistical significance measure for each motif as defined in (expressed as integer (e.g. -10), equivalent to 1 x 10^E-value^ with lower E-values being more statistically significant), with a motif overall E-value significance cutoff of -02 as recommended in MEME, with the exception of motif 29 which had an E-value = 1; and each motif’s MEME-identified amino acid consensus sequence (LOGO) as determined (within MEME) using the sequence display program LOGO. Motif E-values are calculated in MEME based on combined motif p-values on individual taxa, with only significant motif p-values (≤1 x 10^-10^) on individual taxa used to calculate E-values. **(C)** Schematic of individual motifs from S1B arranged from left to right in order of increasing E-values (from most to least significant). Widths of individual motifs are displayed to scale for motifs having 16 aa or greater; motif widths of 8 - 15 aa are displayed as equal to 16 aa for visibility. Each motif is color-coded according to the motif group to which it is assigned. Orange bar indicates E-value = -02 statistical significance cutoff. Scale bar = 50 aa (with the exceptions stated above).

**Fig. S2.**
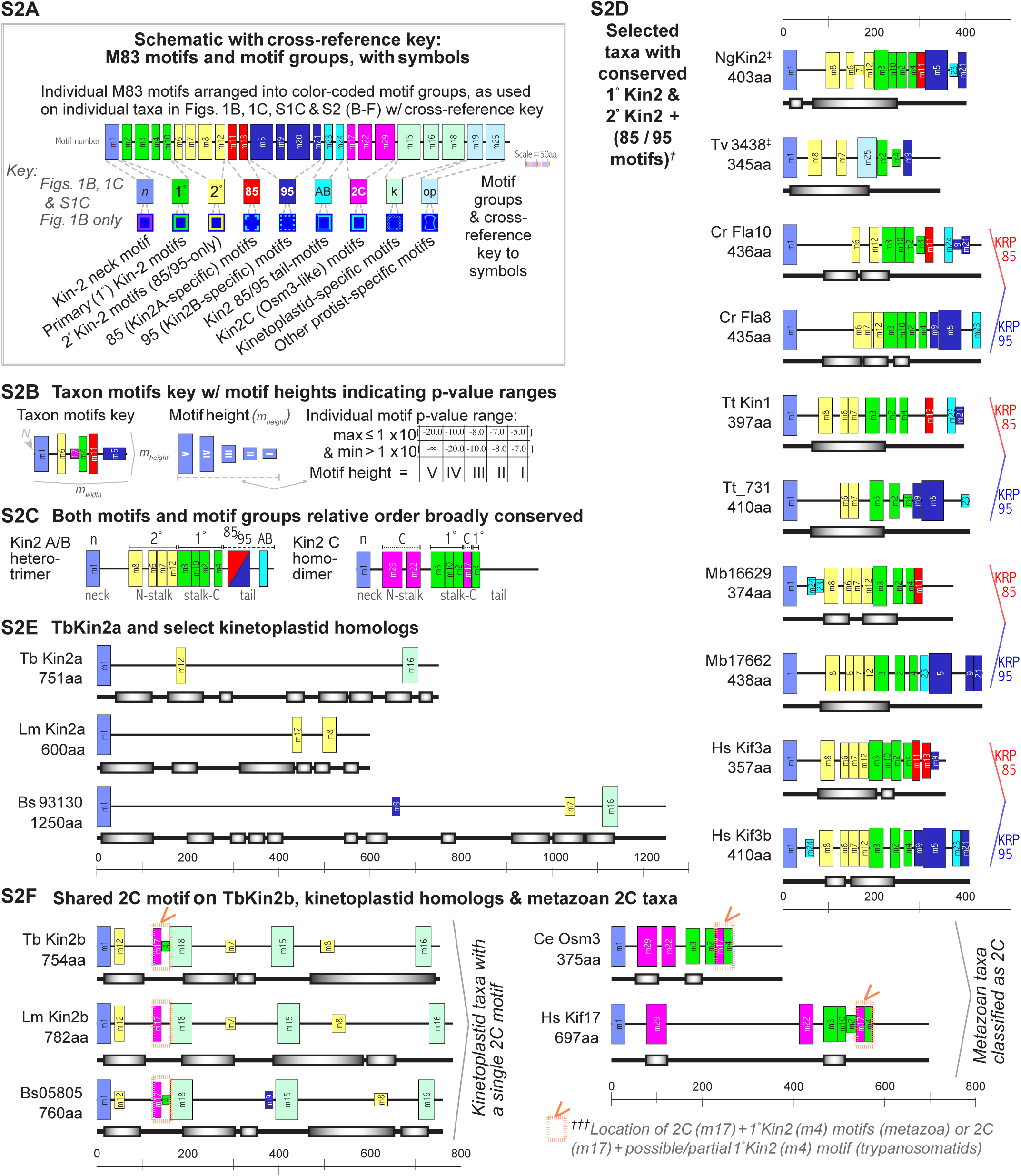
Kinesin-2 NST motifs and motif groups in individual taxa. **(A)** Schematic of individual motifs arranged into motif groups. Motif groups are cross-referenced to motif group symbols that are used in Fig. 1 (B-D). In Fig. 1 and Fig. S2, references to motif group 2A/ KRP85/ 85 are equivalent, as are references to motif group 2B/ KRP95/ 95, and motif group 2C/ OSM3. See Fig. S1C legend for motif scaling information. **(B)** Key showing how the height of motif boxes on individual taxa scales with motif p-value ranges. Roman numerals from V to I show taxon-specific individual motif p-value ranges as follows: V = p-value ≤ 1 × 10^-20^, IV = p-value >1 × 10^-20^ and ≤1 × 10^-10^, III = p-value >1 × 10^-10^ and ≤1 × 10^-08^, II = p-value >1 × 10^-08^ and ≤1 × 10^-07^, I = p-value >1 × 10^-07^ and ≤1 × 10^-05^. **(C)** Schematic showing that motif order is generally conserved among kinesin-2 proteins as depicted. **(D)** NST motifs for a selected group of heterotrimeric kinesin-2A and -2B proteins across a broad evolutionary backdrop illustrate sequence motif conservation noted in (C). In particular, we observed a high frequency of 1° motifs 2, 3, 4 on taxa, especially signature residues [F - I - P] that terminate 1° motif 2, and [W/Y - 6X (with 1-3 E/D) - W] in 1° motif 4, which were observed previously. **(E)** NST motifs for TbKin2a and homologs LmKin2a and Bs93130. One common kinesin-2 motif (2° motif 12) had a significant p-value on 2 of 3 taxa. **(F)** NST motifs for TbKin2b, the related LmKin2b and Bs05805, as well as the kinesin-2C proteins CeOSM3 and HsKif17. Note that 2C motif 17 is followed directly by 1° motif 4 for both metazoan taxa, 1° motif 4 for *T. brucei* and *B. saltans* taxa and a possible (low p-value) amino acid signature for *L. major* (not shown). TbKin2b and kinetoplastid homologs also have a predicted 2° motif 12, here located consistently just after neck domain of the 3 kinetoplastid proteins.

**Fig. S3.**
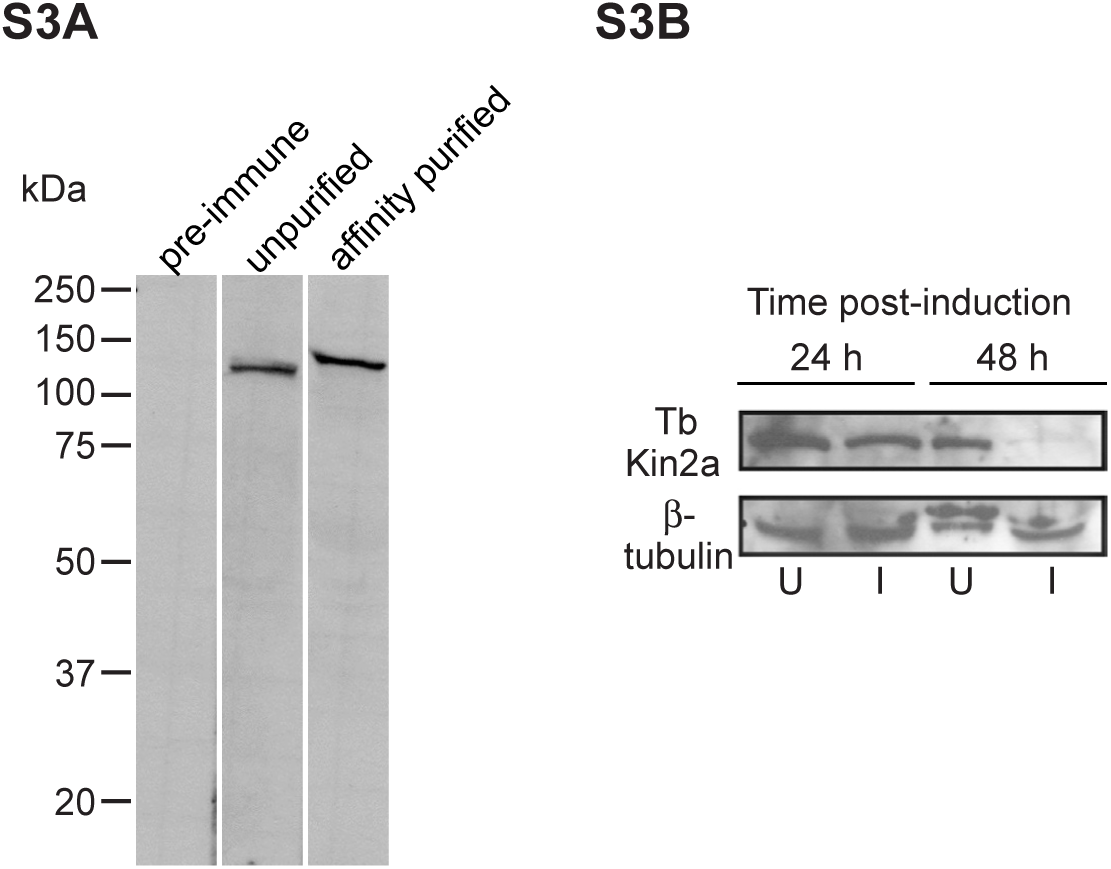
**(A)** Immunoblots of whole-cell extracts from *T. brucei* probed with preimmune serum (left), unpurified post-immune serum (center), and affinity-purified polyclonal antibody (right). **(B)** Immunoblots of whole-cell extracts probed with anti-TbKin2a antibody (top) and anti-β-tubulin antibody KMX (bottom), from uninduced cells (U), or TbKin2a RNAi induced cells (I) at 24 and 48 h post induction.

